# Nanoscale volumetric fluorescence imaging via photochemical sectioning

**DOI:** 10.1101/2024.08.01.605857

**Authors:** Wei Wang, Xiongtao Ruan, Gaoxiang Liu, Daniel E. Milkie, Wenping Li, Eric Betzig, Srigokul Upadhyayula, Ruixuan Gao

## Abstract

Optical nanoscopy of intact biological specimens has been transformed by recent advancements in hydrogel-based tissue clearing and expansion, enabling the imaging of cellular and subcellular structures with molecular contrast. However, existing high-resolution fluorescence microscopes have limited imaging depth, which prevents the study of whole-mount specimens without physical sectioning. To address this challenge, we developed “photochemical sectioning,” a spatially precise, light-based sample sectioning process. By combining photochemical sectioning with volumetric lattice light-sheet imaging and petabyte-scale computation, we imaged and reconstructed axons and myelination sheaths across entire mouse olfactory bulbs at nanoscale resolution. An olfactory-bulb-wide analysis of myelinated and unmyelinated axons revealed distinctive patterns of axon degeneration and de-/dysmyelination in the neurodegenerative mouse, highlighting the potential for peta- to exabyte-scale super-resolution studies using this approach.

## Main Text

Deciphering the brain to understand human cognition, behavior, and consciousness is one of humanity’s most complex and significant undertakings. This endeavor has catalyzed recent development of key technologies to uncover the anatomical complexity and molecular inner workings of the brain. Among these techniques, tissue clearing and expansion has emerged as an effective tool for optical investigation of intact brain tissue (*1–3*). In particular, hydrogel-based tissue clearing and expansion methods have enabled spatial mapping of biomolecules with nanoscale resolution and minimized optical scattering/aberration when combined with fluorescence volumetric imaging (*4–6*). For example, the combination of tissue expansion and light-sheet fluorescence microscopy has created an optimal platform to study subcellular structures and molecular compositions of invertebrate and rodent brains, including their synaptic ultrastructure and structural connectivity, with high imaging speed, low photobleaching rate, and 3D nanoscale resolution (*7–9*).

For nanoscale imaging of expanded tissue, both the expansion factor and the imaging setup contribute to the overall imaging resolution and scalability. Optimizing both is crucial for achieving isotropic resolution, effective optical sectioning, high sensitivity, low photobleaching, and high imaging speed in a large expanded sample. In expansion lattice light-sheet microscopy, for example, the setup offers imaging speed exceeding 100 million voxels per second while providing sub-100 nm resolution and minimal photobleaching at ∼4-fold expansion (*7*). However, the maximum imaging depth for the expanded sample is physically limited by the inherent trade-off between the sample dimensions, objective numerical aperture (NA), and working distance (*10*). Recent studies have adopted ultra-long-working-distance objectives for large-scale imaging (*11*). Nevertheless, balancing NA and working distance continues to pose an obstacle in scaling up existing expansion-based nanoscale imaging methods to whole-mount specimens without sample sectioning.

Recent advances in electron and fluorescence microscopy offer several best practices for sample-sectioning-based volumetric imaging. Techniques such as serial-section EM and focused-ion-beam (FIB)-SEM employ serial sectioning or surface-ablation of the sample followed by on-section or block-face imaging (**Fig. 1A**) (*12–15*). Large volumetric fluorescence imaging of fixed or cleared samples is achieved through sequential physical sectioning followed by on-section or on-block volumetric imaging (*16–21*). However, applying such physical sectioning processes to hydrogel-embedded specimens can lead to sample distortion, tearing, and loss (**fig. S1, fig. S2, Movie S1**). In addition, existing vibratomes and (ultra)microtomes struggle to accommodate the increased size of mammalian tissues, especially of those expanded (*22*). Even with a robust physical sectioning and imaging pipeline, the sample loss and distortion from the sectioning process pose computational challenges when a large sample volume needs to be stitched precisely at high resolution (**fig. S1**) (*22*, *23*). Combined, these factors drastically restrict the dimensions and shapes of samples that can be practically imaged using conventional nanoscale fluorescence imaging techniques.

**Fig. 1.**
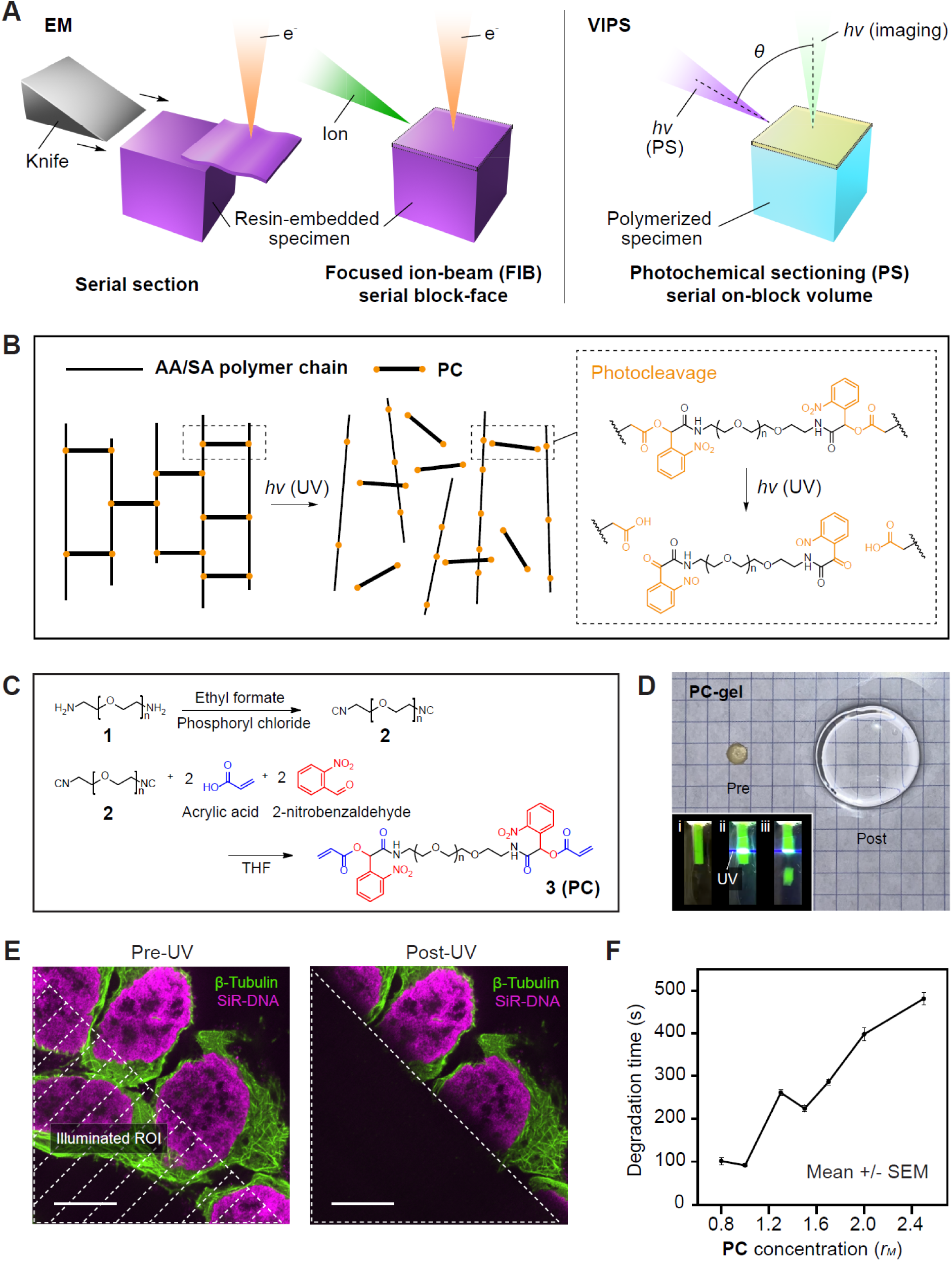
Volumetric imaging of biological specimens via photochemical sectioning (VIPS). (**A**) Schematics showing volumetric imaging of biological specimens via serial section and focused-ion-beam (FIB) serial block-face electron microscopy (EM, left), and via photochemical sectioning (PS) serial on-block volumetric optical imaging (VIPS, right). Depending on the optical setup and configuration, the angle (*θ*) between the imaging [*hv* (imaging)] and the photochemical sectioning [*hv* (PS)] light beams ranges from 0 to 90 degrees in VIPS. (**B**) Schematics showing photochemical degradation (photodegradation) of acrylamide (AA)/sodium acrylate (SA) polymer chains via photocleavage of the crosslinker (photocleavable crosslinker, PC). Inset, schematics showing photocleavage of PC under UV illumination [*hv* (UV)]. (**C**) Synthesis of PC (compound **3**). The second step employs a Passerini reaction for facile synthesis. (**D**) Pre- and post-expansion images of PC-crosslinked polyacrylamide/sodium polyacrylate hydrogel (PC-gel). Grid size: 5 mm. Inset, camera images showing the same piece of PC-gel (fluorescently labeled with a green dye) in water before (i), during (ii), and after (iii) UV (405 nm) laser sheet illumination from the side. (**E**) Photodegradation of HEK cells, fluorescently labeled for microtubules (β-tubulin, AF488) and nuclei (SiR-DNA) and expanded with the PC-gel. The shaded area in the pre-UV image indicates the region of interest (ROI) where the UV (405 nm) laser was illuminated. Scale bars: 10 µm (48 µm). Here and after, unless otherwise noted, scale bars are provided at pre-expansion scale (with the corresponding post-expansion size indicated in brackets). (**F**) Photodegradation time of PC-gel-expanded HEK cells under the same UV (405 nm) laser illumination with different PC concentrations [error bar, standard error of the mean (SEM); n = 3 gels]. *rM*, molar ratio of PC to bis-acrylamide, which is used in the non-photodegradable, bis-acrylamide-crosslinked polyacrylamide/sodium polyacrylate gel (Bis-gel).

To overcome the challenges of traditional imaging and sectioning techniques, we introduce <u>V</u>olumetric <u>I</u>maging of biological specimens via <u>P</u>hotochemical <u>S</u>ectioning (VIPS), a novel method for nanoscale volumetric fluorescence imaging. VIPS employs a spatially precise photochemical degradation process to perform sequential etching and on-block volumetric imaging of biological specimens (**Fig. 1A, Movie S2**). We termed this light-based sample sectioning process “photochemical sectioning” to distinguish it from conventional physical sectioning methods. To implement VIPS, we first designed a photodegradable superabsorbent hydrogel that degrades rapidly and completely under UV illumination. We then confirmed that fluorescently labeled tissue embedded in this hydrogel could be photodegraded in a spatially confined manner using both single- and multi-photon illumination. Next, by combining sequential on-block volumetric lattice light-sheet imaging, light-sheet photochemical sectioning, and petabyte-scale computation, we imaged and reconstructed axonal projections and myelination sheaths across two complete mouse olfactory bulbs at nanoscale resolution. Finally, we performed a comparative analysis of the wild-type and neurodegenerative mouse olfactory bulb, which revealed unique spatial patterns of axon degeneration and de-/dysmyelination in the neurodegenerative sample at single-axon resolution.

### Design and synthesis of photodegradable hydrogel

Photoresponsive hydrogels have been used in multiple studies to modulate mechanical and chemical environments around encapsulated cells (*24–27*). After evaluating various photochemistries for their optical properties and compatibilities with tissue expansion and clearing, we selected polyacrylamide hydrogel for VIPS. This polymer, widely used for tissue clearing and expansion, can undergo swift and complete photodegradation by incorporating a photocleavable crosslinker (PC) to its polymer network (**Fig. 1B**). To enhance the usability and accessibility of VIPS, we prioritized the following design principles for PC: (a) a photocleavage wavelength distinct from popular fluorophores; (b) bio-orthogonal and water-soluble; (c) a low molecular weight for enhanced sample permeability; (d) synthesizable in a minimal number of steps with common wet-lab equipment.

Following these principles, we designed the molecular structure and synthetic pathway of PC (**Fig 1C**). This PC molecule, equipped with a ∼2 kDa polyethylene glycol (PEG) backbone and two photocleavable *o*-nitrobenzyl moieties, can be synthesized in the ambient environment using standard glassware and wet-lab equipment (**Materials and Methods**, **fig. S3**). Using PC as the crosslinker, we then synthesized a photodegradable version of the polyacrylamide/sodium polyacrylate hydrogel via free-radical chain-growth polymerization (**Fig. 1D**). The resulting optically transparent and mechanically elastic hydrogel (“PC-gel”) underwent the same osmotic swelling as its non-photodegradable counterpart (“Bis-gel”, polyacrylamide/sodium polyacrylate hydrogel crosslinked with bis-acrylamide) (*4*). Consistent with the Bis-gel, the expansion factor of the PC-gel increased as the PC concentration decreased during polymerization (**fig. S4**) (*4*). In practice, ∼4-5-fold expanded sample was the most convenient to handle, as the PC-gel maintained a high mechanical integrity around these expansion factors. Therefore, unless otherwise noted, we used this expansion factor for all biological sample preparations. Lastly, we measured global expansion isotropy of the PC-gel and found its ∼1-5% root-mean-square (r.m.s.) error comparable to that of the Bis-gel (**fig. S5**) (*4*).

### Spatially precise photodegradation of PC-gel

Under a continuous illumination of a 365 nm light, the bulk PC-gel underwent complete photodegradation, transforming into a disintegrated liquid state (**fig. S6**). Although PC photocleavage occurs most efficiently around 270-350 nm, the commonly used 405 nm light is also effective since PC absorbs at this wavelength (*24*). We first validated that such photodegradation can be performed in a spatially precise manner by illuminating a piece of PC-gel from the side with a thin sheet of 405 nm laser [**Fig. 1D** **(inset), Movie S3**]. This spatially targeted approach further enabled single-photon photodegradation of PC-gel-expanded cell samples across a defined region-of-interest (ROI) on a confocal microscope (**Fig. 1E, Table S1**). The Bis-gel controls confirmed that the signal loss within the photodegraded ROI resulted from the diffusion of freed, originally gel-anchored fluorophores, rather than photobleaching (**fig. S7**, **fig. S8**). Finally, we measured the PC-gel photodegradation time at varying PC concentrations using the same cell assay (**Fig. 1F**) (*4*). As a result, PC-gels with higher PC density required longer illumination times at the same laser power to complete the photodegradation.

### Nanoscale volumetric fluorescence imaging via two-photon photochemical sectioning

In VIPS, PC-gel-embedded samples go through iterative rounds of on-block volumetric imaging and photochemical sectioning while maintaining a sufficient overlap between each imaging round. Such overlap provides registration landmarks necessary for accurate stitching of the imaged subvolumes. Single-photon widefield or confocal illumination is thus not suitable for photochemical sectioning in a single-objective microscope setup as its out-of-focus light passes through the PC-gel and causes undesirable photodegradation outside the intended volume. To spatially confine PC-gel photodegradation, we exploited the nonlinear optical properties of two-photon (2P) absorption. In this “2P VIPS” method, we first volumetrically imaged the sample, and then raster-scanned a 2P-focused spot to photodegrade the top portion of the sample before iterating this process to reach an imaging depth beyond the objective working distance (**Fig. 2A**). During this process, we ensured the degraded volume was smaller than the imaged volume to maintain sufficient overlaps between the subvolumes.

**Fig. 2.**
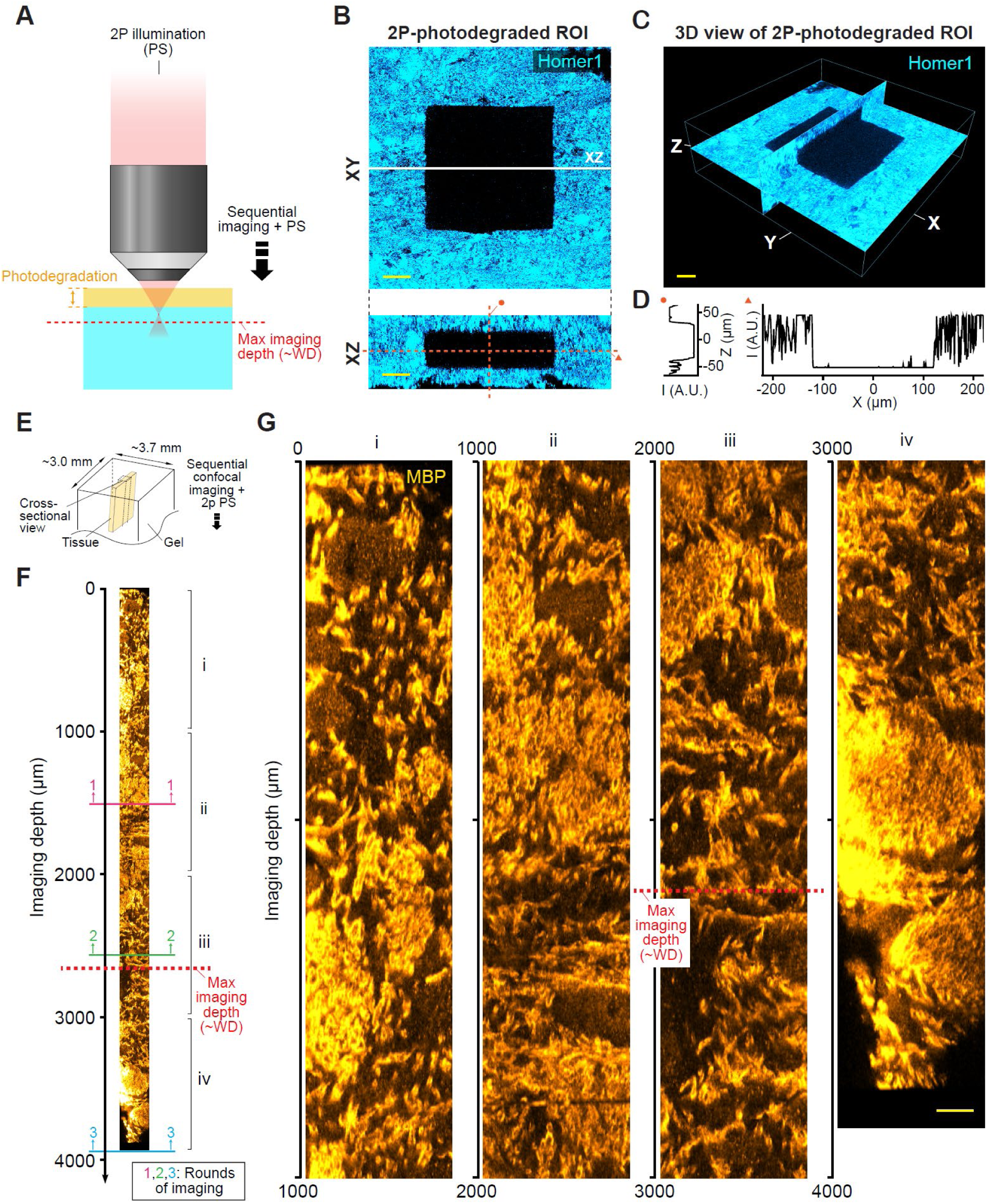
Nanoscale volumetric fluorescence imaging of mouse brain tissue with two-photon (2P) VIPS. (**A**) Schematics showing an exemplary optical setup for 2P VIPS. Imaging beyond the microscope working distance (WD) is achieved by sequential volumetric fluorescence imaging and 2P photochemical sectioning (PS). (**B**) XY (top) and XZ (bottom) views of a 2P-photodegraded ROI in an expanded mouse brain tissue sample. The brain tissue was fluorescently labeled for synaptic proteins (Homer1) and expanded with the PC-gel. Scale bars: 50 µm (post-expansion scale). (**C**) 3D rendered view of the 2P-photodegraded ROI in **B**. (**D**) Line profiles of fluorescence intensity along the orange dotted lines in **B**. (**E**) Schematics showing the block-face of a mouse brain tissue strip vertically embedded and expanded in the PC-gel and imaged with 2P VIPS. (**F**) Vertical cross-sectional view of the PC-gel-expanded mouse brain tissue strip imaged across a ∼3.8 mm imaging depth (beyond the ∼2.6 mm objective WD) after three rounds of sequential on-block volumetric confocal imaging and 2P photochemical sectioning (with no sectioning after the last round of imaging). The tissue was fluorescently labeled for myelin sheaths (myelin basic protein, MBP). The numbered (1–3) lines and arrows indicate the maximum imaging depth reached after each round of imaging. The three imaged subvolumes were computationally stitched using the 3D rigid registration pipeline. (**G**) Enlarged vertical cross-sectional view of the imaged tissue strip from regions i-iv in **F**. Scale bar: 50 µm (post-expansion scale).

To test the spatial control of 2P photochemical sectioning, we used a 740 nm 2P laser to photodegrade a 3D ROI within a PC-gel-expanded mouse tissue sample densely labeled for synaptic proteins (**Fig. 2B**) (*24*). After photodegradation, the illuminated region showed both lateral and axial confinements that matched well with the defined 3D ROI (**Fig. 2C**-**2D**). Once validated, we applied 2P VIPS to a PC-gel-expanded mouse brain slice to overcome the working distance limit of a commercially available objective (**Fig. 2E**). We performed three rounds of on-block volumetric confocal imaging with depths of ∼1.5 mm, ∼1.5 mm, and ∼1.8 mm, respectively, with each imaging round, except for the last, followed by a ∼1.0 mm 2P photodegradation step across the entire sample block-face. As a result, the three stitched subvolumes spanned a ∼3.8 mm thick volume from the original sample surface, far beyond the objective’s working distance of ∼2.6 mm (**Fig. 2F-2G**).

### Nanoscale volumetric fluorescence imaging via light-sheet photochemical sectioning

While volumetric confocal imaging and 2P photochemical sectioning effectively extends the imaging depth, its relatively slow imaging speed and susceptibility to photobleaching restricts its use for large-volume imaging. Light-sheet offers enhanced imaging speed and reduced photobleaching, which makes it an ideal platform for VIPS (*7*, *8*). Hence, we implemented “light-sheet VIPS” with sequential volumetric light-sheet imaging and light-sheet photochemical sectioning (**Fig. 3A, Movie S2**).

**Fig. 3.**
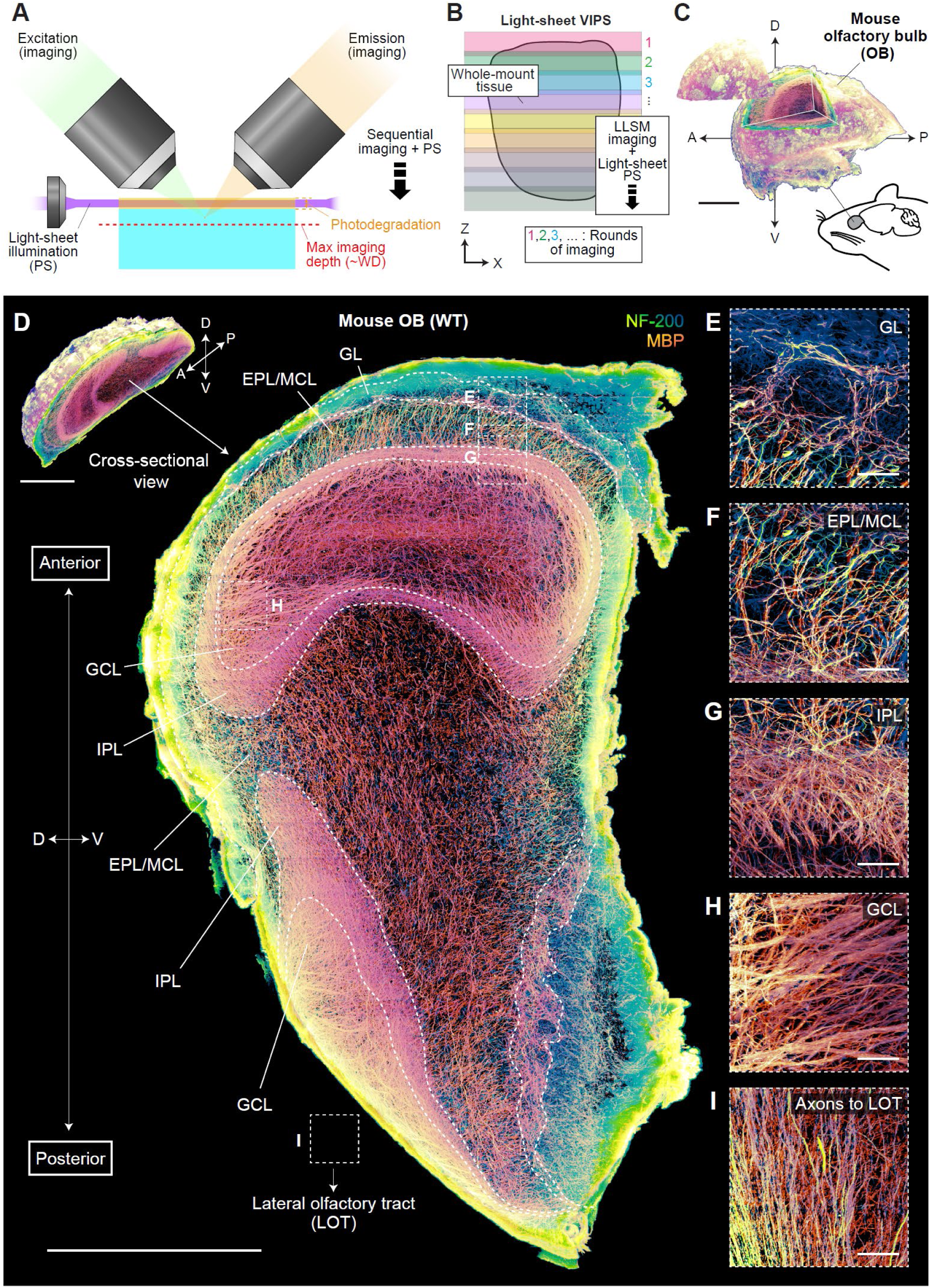
Nanoscale volumetric fluorescence imaging of mouse olfactory bulb (OB) with light-sheet VIPS. (**A**) Schematics showing an exemplary optical setup for light-sheet VIPS. Imaging beyond the microscope WD is achieved by sequential volumetric fluorescence imaging and light-sheet photochemical sectioning (PS). The top excitation and emission objectives show one possible optical configuration for the imaging. (**B**) Schematics showing imaging of expanded whole-mount tissue with sequential lattice light-sheet microscopy (LLSM) imaging and light-sheet photochemical sectioning. Volumetric overlaps between the imaged subvolumes are maintained for lossless reconstruction of the entire imaged volume. (**C**) (Top) 3D rendered view of a 7-week wild-type (WT) mouse OB (with a wedge of the volume moved to the side) imaged using sequential lattice light-sheet imaging and light-sheet photochemical sectioning. The tissue was fluorescently labeled for axons (NF-200) and myelin sheaths (MBP). The dataset was downsampled 5-fold in each direction for visualization. (Bottom) The approximate location of OB in the mouse brain. Scale bar: 1 mm (1.93 mm). A: anterior; P: posterior; D: dorsal; V: ventral. (**D**) 3D rendered slab view (∼80 µm thick at pre-expansion scale) of the imaged WT mouse OB. The dataset was downsampled 5-fold in each direction for visualization. Scale bar: 1 mm (1.93 mm). Top inset, 3D rendered view of a portion of the wild-type mouse OB, exposing the same slab view. Scale bar: 1 mm (1.93 mm). (**E-I**) Full-resolution maximum intensity projection (MIP) view (∼80 µm thick at pre-expansion scale) of the boxed regions in **D** near GL (**E**), EPL/MCL (**F**), IPL (**G**), GCL (**H**), and LOT (**I**). Region **I** is from a different slab view from **D**. Scale bars: 50 µm (96.5 µm). GL: glomerular layer; EPL: external plexiform layer; MCL: mitral cell layer; IPL: internal plexiform layer; GCL: granule cell layer; LOT: lateral olfactory tract.

As an initial validation, we tested whether light-sheet VIPS could extend the imaging capacity of expansion lattice light-sheet microscopy to thick samples beyond the microscope’s working distance (*7*). To achieve spatially precise photodegradation parallel to the sample surface, we integrated a separate UV light-sheet illumination path (**fig. S9**). This implementation generates a thin UV light-sheet compatible with both single and dual objective configurations since it bypasses the objectives and arrives parallel to the sample surface. Characterization using a 100 mW 405 nm laser revealed that a complete photodegradation of the PC-gel-embedded samples could be completed in ∼20 minutes for ∼4-5-fold expansion in water and in ∼2 hours for ∼2-fold expansion in 1x PBS (**Materials and Methods**).

We then validated the capabilities of light-sheet VIPS on a ∼4.4-fold expanded human hippocampus tissue sample labeled for axons and myelin sheaths. We performed eight rounds of sequential volumetric lattice light-sheet imaging followed by light-sheet photochemical sectioning across a ∼1.6 mm sample thickness at post-expansion scale—five times the imaging depth achievable without VIPS (**fig. S10**). The imaged subvolumes were then registered and stitched using PetaKit5D with a pre-expansion Nyquist-sampled voxel size of 22 by 22 by 59 nm (*28*). The final dataset allowed us to trace and reconstruct individual axons with diameters as small as ∼150 nm within the imaged volume, confirming that the pipeline preserved the connectivity and traceability of anatomical features across multiple rounds of imaging and photochemical sectioning. These results demonstrate that light-sheet VIPS can be potentially scaled up for high-throughput, super-resolution imaging of whole-mount expanded samples.

### Mapping axons and myeline sheaths in whole mouse olfactory bulb

The mammalian olfactory bulb (OB) is a complex neural structure essential for processing and relaying olfactory information. Understanding its anatomical and molecular architecture can provide critical insights into the anatomy and function of mammalian sensory system. To showcase the scalability of light-sheet VIPS, we performed a two-color imaging of an entire 7-week wild-type mouse OB. For this proof-of-concept experiment, we used ∼2-fold expansion with a pre-expansion Nyquist-sampled voxel size of 56 by 56 by 135 nm, as this resolution is sufficient to trace and reconstruct most axons in the mouse brain (*29*). Opting for higher expansion factors would increase the data volume to tens of petabytes for the raw, intermediate, and fully processed volumes, easily surpassing our currently accessible data handling and computation capabilities.

After mounting the expanded OB, we imaged the entire sample volume (∼9.4 by 6.4 by 4.3 mm at post-expansion scale) with the peak acquisition rates approaching 2 teravoxels per hour. The OB imaging was completed in 10 days, utilizing nine rounds of interleaved on-block volumetric lattice light-sheet imaging and light-sheet photochemical sectioning, totaling ∼142 hours of imaging and ∼20 hours of photodegradation (**Fig. 3B**). This process yielded 17,280 image tiles, constituting ∼0.5 petabytes of raw data, which were computationally combined to reconstruct the entire mouse OB (**Fig. 3C, Movie S4**). To process this petabyte-scale image dataset with academic-scale computing resources, we built our pipelines using PetaKit5D, a high-performance computing framework designed for efficient image reading, writing, geometric transformations, stitching, and deconvolution (**Materials and Methods**) (*28*). Using this pipeline, we were able to deskew, rotate, deconvolve, and stitch the entire imaged volume with drastically reduced computing time. Moreover, the gentle and spatially precise nature of photochemical sectioning was crucial for achieving lossless overlap when stitching together the imaged subvolumes.

The reconstructed wild-type mouse OB dataset revealed distinct axonal projections and myelination patterns across all its major anatomical layers (**Fig. 3D**). Detailed 3D visualization highlighted unique axonal and myelination patterns within individual glomeruli of the glomerular layer (GL) (**Fig. 3E**). The vertical axonal projections of tufted cells traversed through the external plexiform layer (EPL) and mitral cell layer (MCL) (**Fig. 3F**), while the dense lateral axonal projections spanned the internal plexiform layer (IPL) (**Fig. 3G**). The axonal bundles from the tufted and mitral cells were observed passing through the granule cell layer (GCL) (**Fig. 3H**), exiting the olfactory bulb, and projecting through the lateral olfactory tract to the olfactory cortex (**Fig. 3I**). Furthermore, the nanoscale resolution of the dataset enabled segmentation, skeletonization, and reconstruction of individual myelinated and unmyelinated axon fibers, allowing for precise measurements such as calculating their total lengths throughout the entirety of mouse OB (**Fig. 4A-4B**, **fig. S11**).

**Fig. 4.**
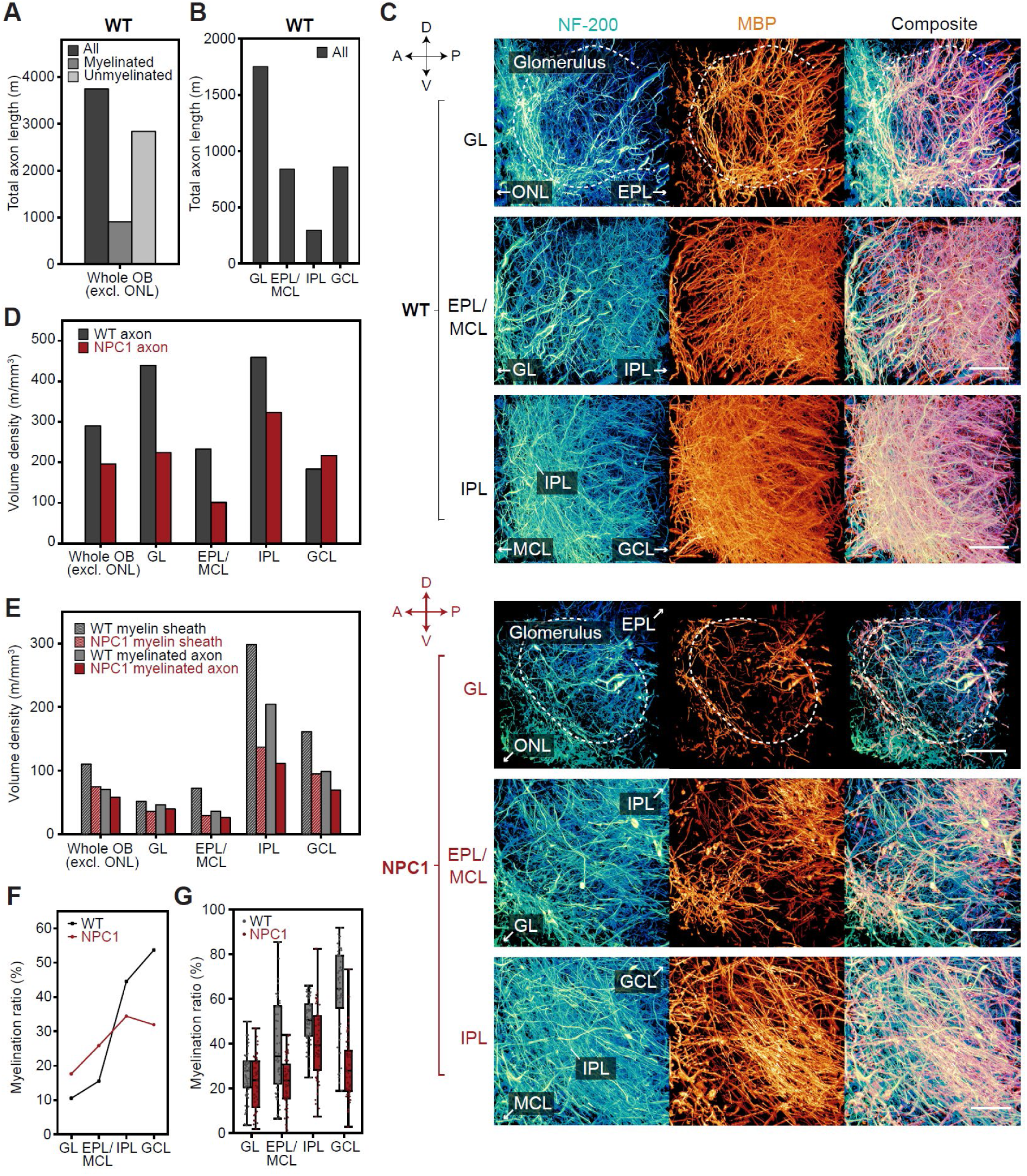
Olfactory-bulb (OB)-wide analysis of axons and myelin sheaths at single-axon resolution. (**A**) Estimated total length of axons, myelinated axons, and unmyelinated axons in wild-type (WT) mouse OB. Here and after, unless otherwise noted, ONL is excluded from the OB-wide analysis. (**B**) Estimated total length of axons in distinct anatomical layers of WT mouse OB. (**C**) Full-resolution 3D rendered view of representative regions in GL, EPL/MCL, and IPL of 7-week WT (top) and NPC1 (*Npc1*^−/−^) (bottom) mouse OBs. NF-200, MBP, and composite views of the same region are shown in separate columns. Scale bars, 50 µm (96.5 µm). A: anterior; P: posterior; D: dorsal; V: ventral. (**D**) Volume density of axons across the entirety and distinct anatomical layers of the WT and NPC1 mouse OB. (**E**) Volume density of myelin sheaths and myelinated axons across the entirety and distinct anatomical layers of the WT and NPC1 mouse OB. (**F**) Myelination ratio of axons across distinct anatomical layers of the WT and NPC1 mouse OB. (**G**) Myelination ratio distribution of 50 ROIs (∼66 by 66 by 87 µm at pre-expansion scale) randomly sampled across distinct anatomical layers of the WT and NPC1 mouse OB. Data are presented as box plots, where the ends of the whiskers represent the maximum and minimum values of the distribution, the dots represent individual values, and the upper, middle, lower line of the box represents the 75th percentile, 50th percentile (median), and the 25th percentile, respectively. ONL: olfactory nerve layer; GL: glomerular layer; EPL: external plexiform layer; MCL: mitral cell layer; IPL: internal plexiform layer; GCL: granule cell layer.

### Axon degeneration and de-/dysmyelination in neurodegenerative mouse olfactory bulb

Axonal degeneration and de-/dysmyelination, often preceding clinical symptoms, are hallmarks of neurodegeneration and aging. In the fatal neurodegenerative lysosomal storage disorder Niemann-Pick type C1 (NPC1) disease, mutations in the *NPC1* gene cause early-stage dysmyelination (*30*). Anatomical and physiological studies have identified pathophysiological changes in the NPC1 animal’s olfactory system as an early indicator of disease progression (*31*, *32*), which is also characterized by the reduced myelin basic protein (MBP) expression in the OB at the neonate and adolescent stage (*33*). However, the precise mechanism behind de- /dysmyelination remains unclear, particularly regarding whether the reduced expression of MBP in the OB persists into adulthood and results in anatomical loss at the single axon/myelin sheath level. Furthermore, the progression of myelin sheath loss across the distinct layers of OB remains unexplored.

To study axon degeneration and de-/dysmyelination in the NPC1 brain, we used light-sheet VIPS to image a 7-week NPC1 (*Npc1^−/−^*) mouse OB (∼2-fold expanded and extending a volume of ∼11.2 by 6.0 by 3.6 mm) (**Movie S5**). The resulting multi-scale WT and NPC1 mouse OB datasets allowed visualizing the axon and myelin sheath morphologies in the corresponding regions across the entire OB (**Fig. 4C, Movie S4-S5**). Close comparison of the two datasets revealed distinct anatomical changes, including decreased axon and myelin sheath densities in the GL, EPL/MCL, and IPL of the NPC1 OB, suggesting potential axon degeneration and de-/dysmyelination in these layers.

OB-wide analysis of axon fibers showed reduced axon density in the NPC1 OB, consistent with previous studies showing early-onset axonal degeneration across all major parts of the NPC1 brain (**Fig. 4D**) (*34–36*). Unexpectedly, the GCL exhibited a higher axon density for the NPC1 sample, while all other layers showed consistent decreases (**Fig. 4D**). Moreover, the severity of axon degeneration varied spatially across the GL, EPL/MCL, and IPL, with substantial reduction in the GL and the least in the IPL. These results suggest that local activity, connectivity, and the extracellular environment within each OB layer are more likely to be responsible for axonal integrity and survival, given that axonal damage often precedes cell body loss (*35*, *37*).

Additionally, we evaluated the myelination profile of both the WT and NPC1 mouse OBs. A reduction in the density of total myelin sheaths and myelinated axons was evident throughout the NPC1 OB affirming that early-stage dysmyelination persists as the animal maturates (**Fig. 4E**) (*33*). Interestingly, the ratio of myelinated axons to all axons (the myelination ratio) varied across the layers, with decreases in IPL and GCL, but increases in GL and EPL/MCL in the NPC1 OB (**Fig. 4F**). This suggests that the effects of dysmyelination may differentially impact axon degeneration across different OB layers. Such spatial heterogeneity also exists locally within the same layer as the myelination ratio and other axonal/myelination parameters varied drastically across the spatially sampled ROIs within each layer (**Fig. 4G, fig. S12**).

## Discussion

VIPS utilizes light-based photochemical sectioning of hydrogel-embedded samples to overcome imaging limitations in sample size, enabling scalable optical nanoscopy of whole-mount samples. Its integration with expansion lattice light-sheet microscopy pipeline (*7*) led to three orders of magnitude increase in the total imaged volume, effectively removing physical constraints on the imaging of large and diverse specimens. A recently developed high-performance computation pipeline was further leveraged to process, visualize, and analyze whole-mount sample datasets at the petabyte scale. The multi-scale nature of the resulting mouse olfactory bulb datasets, spanning nearly five orders of magnitude from tens of nanometers to millimeters, provided biological insights unattainable by sampling a large number of smaller ROIs across the sample (e.g., **Fig. 4F-4G, fig. S12**). A similar approach can be extended to imaging and analysis of samples with higher resolution and denser labeling. Through iterative high-isotropy expansion, for example, the effective resolution of VIPS can be enhanced to the sub-10 nm scale (*38–41*). When paired with pan-membrane or protein labeling, VIPS can offer a viable avenue for dense reconstruction and synaptic-level connectomics at the whole-brain level (*42*, *43*).

While the initial demonstration of VIPS was promising, several challenges remain. Processing petabyte-scale data using state-of-the art computational pipelines revealed residual distortions at subvolume interfaces due to limitations of the current rigid 3D registration and stitching frameworks. Although these distortions do not disrupt anatomical continuity, the development of non-rigid 3D registration methods at this scale is essential to address non-linear sample-deformation-induced stitching errors. Advances in computational software are expected to tackle this challenge and facilitate seamless stitching of datasets at this scale.

To achieve biologically meaningful comparisons, additional replicates are needed. However, the high costs of computational storage infrastructure pose a barrier and limits the number of replicates for a statistically significant number of mouse OB datasets. As computational hardware advances, these limitations are expected to diminish, enabling the study of larger, whole-mount specimens with a focus on biological stereotypy and variability.

Scaling VIPS to the exabyte level (“Exascale VIPS”) for whole-mount sample imaging at nanoscale resolution is possible, but requires advancements across several areas. These include the development of uniform and scalable staining pipelines to eliminate variations in immunostaining across the specimens. Reliance on immunofluorescence for molecular labeling can introduce biases from off-target, low on-target, and gradient of antibody binding. Such biases can skew downstream analyses, such as the estimation of axon and myelin sheath densities across the mouse OBs. Ongoing effort to improve immunostaining techniques or a transition to endogenous labeling are essential to mitigate these biases (*44–46*). Improvement of imaging hardware such as larger sample stages, imaging chambers to accommodate entire samples, parallelized simultaneous photodegradation and imaging, powerful photochemical sectioning lasers, implementation of non-diffracting 2P light-sheets for more targeted control over the gel degradation, larger frame sCMOS cameras, and simultaneous multi-color imaging are expected to accelerate the data throughput.

The massive datasets produced in a single experiment by these technologies pose challenges in visualization and analysis. Solutions such as NVIDIA Index may provide promising avenues for real-time visualization at the peta- to exabyte scale leveraging recent advances in GPU hardware and scalable software (*28*). However, manually segmenting and quantifying details of subcellular features across peta- to exavoxel-sized volumes is impractical. The advances in artificial intelligence (AI) (*47*), particularly in large multimodal AI models, may offer a path forward. Once available, these models will allow researchers to feed multi-dimensional data and output optimally reconstructed volumes with a database of queryable segmented features and their quantified properties. These outputs, when combined with large language models, enable direct data mining, thus unlocking the full potential of data generated using modern imaging technologies, including VIPS.

## Acknowledgments

We thank J. Zak for discussion on mouse olfactory system and assistance with data analysis. We thank S. Cologna for mouse sample preparation, A. Schantz and D. Keene for human sample collection, J. Sun and W. Cho for cell cultures, Y. Wang for assistance with the macroscale light-sheet photochemical sectioning experiment, and R. Torrez, J. Iwasa, and S. Kwon for assistance with animation and scientific illustration. We thank the Center for Advanced Microscopy & Nikon Imaging Center at Northwestern University and the Chicago Biomedical Consortium (CBC) for access to fluorescence microscopy facilities. We thank the NVIDIA IndeX team for sharing the NVIDIA Index software. We thank John White for managing the ABC computing cluster. We gratefully acknowledge the support of this work by the Laboratory Directed Research and Development (LDRD) Program of Lawrence Berkeley National Laboratory under US Department of Energy contract No. DE-AC02-05CH11231. This research used resources of the National Energy Research Scientific Computing Center (NERSC), a U.S. Department of Energy Office of Science User Facility located at Lawrence Berkeley National Laboratory, operated under Contract No. DE-AC02-05CH11231 using NERSC award DDR-ERCAP0025501.

## Funding

National Institutes of Health grant UG3MH126864 (RG, co-I)

University of Illinois Chicago Startup Fund (RG)

Philomathia Foundation Gift (SU, EB)

Lawrence Berkeley National Laboratory’s LDRD Program grants 7647437 & 7721359 (SU)

HHMI (EB)

Chan Zuckerberg Initiative Imaging Scientist program grants 2019-198142 & 2021-244163 (SU)

Chan Zuckerberg Biohub – San Francisco Investigator gift funding (SU) Sloan Foundation grant G-2022-19390 (SU)

## Author contributions

Conceptualization: RG, SU

Methodology: WW, RG, XR, GL, SU

Software: XR, DM, SU

Formal analysis: XR, SU, GL, WW, RG

Microscope design, construction, and validation: GL, DM, SU, EB

Visualization: SU, RG, XR, WW

Investigation: WW, XR, GL Resources: WL, WW

Data curation: XR, GL, SU

Funding acquisition: SU, EB, RG

Project administration: RG, SU Supervision: RG, SU

Writing – original draft: RG

Writing – review & editing: WW, XR, GL, DM, WL, EB, SU, RG

## Competing interests

RG is a co-inventor of multiple patents related to expansion microscopy. The other authors declare that they have no competing interests. Portions of the technology described herein are filed for patent.

## Data and materials availability

Custom-synthesized materials and reagents used in this manuscript will be made freely available within the capacity of RG lab for researchers under materials transfer agreements (MTAs). Custom scripts used for petabyte-scale image data processing and analysis are available on GitHub at https://github.com/abcucberkeley. The raw data for this paper exceeds a petabyte in total and thus is not practical to be hosted on currently available public repositories. All data used in this manuscript will be made freely available upon request with a feasible data transfer and storage mechanism (such as Globus endpoint, or cloud buckets). All data are available in the main text or the supplementary materials.

## Supplementary Materials

Materials and Methods

Supplementary Text

Figs. S1 to S12

Tables S1

References (*1–50*)

Movies S1 to S5

