## Supplementary Materials for "Nanoscale volumetric fluorescence imaging via photochemical sectioning"

Wei Wang *et al.*

Corresponding authors:
Ruixuan Gao,
Srigokul Upadhyayula,

**The PDF file includes:**

Materials and Methods
Supplementary Text
Figs. S1 to S12
Tables S1
Captions for Movies S1 to S5
References

**Other Supplementary Materials for this manuscript include the following:**

Movies S1 to S5

### Materials and Methods

#### 1. Material Synthesis

Unless otherwise noted, all chemicals and reagents were obtained from Millipore Sigma.

*a. Synthesis of photocleavable crosslinker (PC)*

*a.1 Synthesis of polyethylene glycol bis(2-isocyanoethyl) ether (**2**)*

An ethyl formate (40 mL) solution of polyethylene glycol bis(2-aminoethyl) ether (**1**, Mn ~2000 g/mol, 1 g, ~0.5 mmol, Zhengzhou Alfa Chemical Co., Ltd) was stirred at reflux (80°C) for 48 hours, and dried under reduced pressure using a rotary evaporator (N-1300, EYELA) (with the water bath temperature set at 42°C, here and after, unless otherwise noted). The light-yellow product was then dissolved in anhydrous tetrahydrofuran (THF, 40 mL) before triethylamine ($Et_3N$, 2.8 mL) was added. Next, the solution was chilled and stirred at -70°C, and a THF (2 mL) solution of phosphoryl chloride ($POCl_3$, 0.4 mL, 4.3 mmol) was added to the reaction mixture dropwise over a period of 20 min. The solution was stirred overnight in an ice bath, poured into a saturated sodium carbonate (Themo Fisher) aqueous solution (150 mL, chilled at 4°C), and further stirred at < 20°C for one hour. The THF layer was then evaporated under reduced pressure and the remaining aqueous layer was extracted five times with 200 mL of dichloromethane (DCM, Themo Fisher) each time. Finally, the combined DCM layer was dried under reduced pressure before ethyl acetate (EA, 10 mL) was added to dissolve the crude product. A large amount of petroleum ether (Thermo Fisher) or hexane (Themo Fisher) was added to the EA solution, and the mixture was placed in an ice bath for 30 min. The precipitates were filtered, washed with cold petroleum ether or hexane, and dried under vacuum overnight to yield the light yellow-white solid product (**2**).

*a.2 Synthesis of photocleavable crosslinker (**PC**, **3**)*

The photocleavable crosslinker (**PC**, **3**) was synthesized using a Passerini reaction [see *Chem Sci* **11**, 8224–8230 (2020); *Polymer Bulletin* **76**, 1471–1487 (2019)]. First, compound **2** from the previous step of synthesis (Mn ~2000 g/mol, 0.95 g, ~0.475 mmol), 2-nitrobenzaldehyde (250 mg, 1.50 mmol), and acrylic acid (120 mg, 1.67 mmol) were dissolved in THF (20 mL). The reaction mixture was then stirred for 24 hours at room temperature in the dark. Next, THF was evaporated from the reaction mixture under reduced pressure, and the yellow oily raw product was dissolved in EA (10 mL) for reprecipitation. Briefly, a large amount of petroleum ether or hexane was added to the EA solution, and the mixture was placed in an ice bath for 30 min. The precipitates were filtered, washed with cold petroleum ether or hexane, and dried under vacuum overnight to yield a light yellow-white powder. Finally, the crude product was dissolved in double-distilled water to a concentration of ~25 mg/mL, dialyzed (Spectra/Por 6 Dialysis Membranes 1kD, Repligen) for 12 hours, dried under reduced pressure, and reprecipitated once again using EA and petroleum ether/hexane to yield the final product of a yellow-white powder (**PC**, **3**).

$^1$H NMR (500 MHz, Chloroform-d, δ): 6.74 (s, 2H; (CO)–C**H**–(Ar)O–, **f**), 6.90 (s, 2H; –N**H**–, **e**), 6.51 (d, 2H; CH2= C**H**–CO, **g**), 6.23 (m, 2H; CO–CH = C**H′**H, **h**), 5.97 (m, 2H; CO–CH = CH′**H**, **i**). Also see **fig. S3**.

#### 2. Biological Sample Preparation

Unless otherwise noted, all chemicals and reagents were obtained from Millipore Sigma.

##### *a. Ethics statements*

All procedures involving mice were performed in accordance with the US National Institutes of Health Guide for the Care and Use of Laboratory Animals and approved by the University of Illinois Chicago Animal Care Committee.

Postmortem human specimens were collected with informed consent from the patients in consultation and compliance with the University of Washington School of Medicine Compliance Office and HIPAA.

##### *b. Human embryonic kidney 293 (HEK 293) cells*

HEK 293 cells (Thermo Fisher or ATCC) were cultured on a 12-mm round coverslip to a confluency of 80-90% (*48*). The cells were then fixed with 4% (w/v) paraformaldehyde (PFA) in 1x PBS for 10 min, washed with 1x PBS twice for 5 min each time, and stored in 1x PBS with

0.03% (w/v) sodium azide. For immunostaining, the cells were permeabilized with 0.1% (w/v) Triton X-100 in 1x PBS for 15 min and incubated in the blocking buffer [5% (v/v) normal goat serum (NGS, Jackson ImmunoResearch) and 0.1% (w/v) Triton X-100 in 1x PBS] for 15 min. The permeabilized and blocked cells were then incubated in the primary antibody (rabbit anti-beta-tubulin antibody, ab6046, Abcam; see **Table S1**) solution (1:200 dilution with the blocking buffer) at 4°C for 12 hours. Next, the cells were washed with blocking buffer four times for 5 min each time, and incubated in the secondary antibody (goat Alexa Fluor 488-conjugated anti-rabbit antibody, A11008, Thermo Fisher or goat ATTO 647N-conjugated anti-rabbit antibody, 40839-1ML-F, Millipore Sigma) solution (1:200 dilution with the blocking buffer) at room temperature for 2-3 hours. Finally, the cells were washed with 1x PBS four times for 5 min each time and stored in 1x PBS for the subsequent gelation procedure.

*c. Mouse*

*c.1 Mouse brain slices*

9-week C57BL/6 mice were anesthetized using ketamine hydrochloride injectable solution (100 mg/mL, Covetrus) and AnaSed Injection (xylazine) sterile solution (20 mg/mL, Akorn, Inc) and transcardially perfused with 4% (w/v) PFA in 1x PBS (10 mL). The brains were carefully dissected from the skull, post-fixed with 4% (w/v) PFA in 1x PBS at 4°C for 1 day, and stored in a sucrose buffer (20% sucrose in 1x PBS) at 4°C. The fixed brains were sectioned using a vibratome (VT1200S, Leica) or flash-frozen with dry ice and then sectioned using a microtome (Model 860, AO Scientific Instruments) to ~40 μm coronal slices. All the sectioned brain slices were stored in 1x PBS with 0.03% (w/v) sodium azide at 4°C.

For immunostaining, the ~40 μm brain slices were permeabilized with 0.1% (w/v) Triton X-100 in 1x PBS for 15 min and incubated in the blocking buffer [5% (v/v) NGS and 0.1% (w/v) Triton X-100 in 1x PBS] at room temperature for more than 6 hours. The brain slices were then incubated in the primary antibody (rabbit anti-NF-200, N4142-.2ML, Millipore Sigma and chicken anti-MBP antibody, PA1-10008, Thermo Fisher, or rabbit anti-Homer 1, 160003, Synaptic Systems) solution (1:200 dilution with the blocking buffer) at 4°C for 2 days. Next, the brain slices were washed with the blocking buffer four times for 30 min each time, and incubated in the secondary antibody (goat Alexa Fluor 488-conjugated anti-rabbit antibody, A11008, Thermo Fisher and goat Alexa Fluor 568-conjugated anti-chicken antibody, A11011, Thermo Fisher, or goat ATTO 647N-conjugated anti-rabbit antibody, 40839-1ML-F, Millipore Sigma, respectively) solution (1:200 dilution with the blocking buffer) at 4°C for 2 days. Finally, the immunostained mouse brain slices were washed with 1x PBS (or the blocking buffer) four times for 30 min each time and stored in 1x PBS for the subsequent gelation procedure.

*c.2 Mouse olfactory bulbs (OBs)*

Heterozygous Balb/c *Npc*$^{\mathrm{nih}}$ (*Npc1*$^{+/-}$) mice were obtained from Jackson Laboratories (RRID: IMSR JAX:003092) and a breeding colony was maintained. 7-week wild-type Balb/c (*Npc1*$^{+/+}$) ("WT") and mutant (*Npc1*$^{-/-}$) mice ("NPC1"), male, were anesthetized using isoflurane inhalation or an injection of ketamine hydrochloride injectable solution (100 mg/mL, Covetrus) and AnaSed Injection (xylazine) sterile solution (20 mg/mL, Akorn, Inc), and transcardially perfused with 4% (w/v) PFA in 1x PBS (10 mL). The brains were carefully dissected from the skull, post-fixed with 4% (w/v) PFA in 1x PBS at 4°C for 1 day, and stored in a sucrose buffer (20% sucrose in 1x PBS) at 4°C. The olfactory bulbs (OBs) were dissected from the rest of the brain and stored in the sucrose buffer at 4°C.

The dissected mouse OBs were immunostained using a modified iDISCO+ protocol as detailed on the continuously-updated website http://idisco.info. First, the OBs were washed with 1x PBS four times for 30 min each time, dehydrated using a methanol (Thermo Fisher)/1x PBS gradient (from 0% methanol, 20% methanol, 40% methanol, 60% methanol, 80% methanol to 100% methanol, 1 hour incubation in each solution), and incubated in methanol at 4°C overnight. The 100% methanol solvent was then replaced with a 66% DCM/33% methanol solution. Next, the OBs were incubated in the 66% DCM/33% methanol solution at room temperature overnight with shaking, washed with 100% methanol twice, and rehydrated with a methanol/1x PBS gradient (from 100% methanol, 80% methanol, 60% methanol, 40% methanol, 20% methanol, to 0% methanol, 1 hour incubation in each solution) at room temperature. Subsequently, the OBs were washed with PTx.2 buffer [0.2% (v/v) Triton X-100 in 1x PBS] at room temperature twice for 1 hour each time.

The pretreated OBs were then permeabilized with a permeabilization buffer [80% (v/v) PTx.2, 20% (v/v) DMSO, 2.3% (w/v) Glycine] at 37°C for 36 hours, and incubated in a blocking buffer [84% (v/v) PTx.2, 6% (v/v) NGS, 10% (v/v) DMSO] at 37°C for 36 hours. Next, the OBs were incubated in the primary antibody (chicken anti-MBP antibody, PA1-10008, Thermo Fisher and rabbit anti-NF-200, N4142-.2ML, Millipore Sigma) solution [1:100 dilution with the staining buffer, which consists of 92% (v/v) PTwH buffer, 3% (v/v) NGS, and 5% (v/v) DMSO] at 37°C for 3 days. The PTwH buffer [1x PBS, 0.2% (v/v) Tween-20, 0.001% (w/v) Heparin] was prepared in advance. The primary antibody solution was then replaced with a freshly prepared primary antibody solution, and the OBs were further incubated at 37°C for 3 days. Finally, the OBs were washed with the PTwH buffer four to five times for 1 hour each time (with the last washing being overnight), and incubated in the secondary antibody (goat Alexa Fluor 488-conjugated anti-rabbit antibody, A11008, Thermo Fisher and goat Alexa Fluor 568-conjugated anti-chicken antibody, A11011, Thermo Fisher) solution (1:100 dilution with the staining buffer) at 37°C for 3 days. The secondary antibody solution was replaced with a freshly prepared secondary antibody solution, and the OBs were further incubated at 37°C for 3 days. The immunostained OBs were washed with the PTwH buffer four times for 1 hour each time and stored in 1x PBS for the subsequent gelation procedure.

*d. Human*

Postmortem human hippocampus specimens were obtained from the University of Washington BioRepository and Integrated Neuropathology (BRaIN) laboratory and the University of Washington Alzheimer's Disease Research Center (ADRC) Precision Neuropathology Core. Briefly, a fresh piece of posterior hippocampus was dissected at rapid autopsy (postmortem interval <12 hours) and immediately fixed in 4% (w/v) PFA in 1x PBS at room temperature for 48 hours. The fixed tissue block was sectioned to ~300-500 μm slices using a vibratome (VT1200S, Leica) and stored in 1x PBS with 0.03% (w/v) sodium azide at 4°C.

The human hippocampus slices were immunostained using the same protocol as the mouse OBs with minor modifications. After primary and secondary antibody staining, lectin staining was performed at 37°C for 3 days using Lycopersicon esculentum (Tomato) lectin (L-1170-2, Vector Laboratories) conjugated with SeTau-647-NHS (K9-4149, SETA BioMedicals) (1:300 dilution with the staining buffer). The stained slices were washed with the PTwH buffer twice for 1 hour each time and stored in 1x PBS for the subsequent gelation procedure.

#### 3. Sample Gelation, Digestion, and Expansion

Unless otherwise noted, chemicals and reagents were obtained from Millipore Sigma. Purified water was obtained from a Milli-Q IQ 7000 Ultrapure Water System (Millipore Sigma).

##### *a. Basic protocol*

Unless otherwise noted, photodegradable samples were prepared using a custom protein-retention expansion microscopy protocol with the photocleavable crosslinker (**PC**, **3**). Briefly, fixed and immunostained tissue samples were incubated in a diluted AcX solution (0.1 mg/mL in 1x PBS) at room temperature overnight. The samples were then washed with 1x PBS twice for 15 min each time and incubated in the monomer solution [1x PBS, 2 M NaCl, 8.625% (w/v) sodium acrylate (Pfaltz&Bauer), 2.5% (w/v) acrylamide, 4.980% (w/v) **PC**] at 4°C overnight. Concentrated stock solutions of ammonium persulfate (APS) [10% (w/v)], tetramethylethylenediamine (TEMED) [10% (w/v)], and 4-hydroxy-2,2,6,6-tetramethylpiperidin-1-oxyl (4HT) [0.5% (w/v)] were added to the monomer solution to a final concentration of 0.2% (w/v), 0.2% (w/v), and 0.01% (w/v), respectively, to yield the gelling solution. Samples in the monomer solution were immediately incubated in the mixed gelling solution at 0°C for 45 min, transferred to a gelation chamber of various shapes and sizes, and gelled in a humidified 37°C incubator for 2 hours ("PC-gel"). The gelled samples were trimmed and immersed in a digestion buffer [1 mM EDTA, 0.5% Triton X-100, 1 M NaCl, 8 units/mL Proteinase K (proK, New England Biolabs) in 1x PBS, no Tris] at room temperature overnight. The digested samples were stored in 1x PBS at 4°C. The stored samples were expanded in purified water three times for 20 min each time before imaging and photodegradation.

Non-photodegradable samples were prepared using the standard protein-retention expansion microscopy protocol with bis-acrylamide as the crosslinker ("Bis-gel") (*48*).

*b. Cell cultures*

Photodegradable cell samples were prepared using a modified basic protocol. The cell samples were directly incubated in a modified gelling solution [1x PBS, 2 M NaCl, 8.625% (w/v) sodium acrylate, 2.5% (w/v) acrylamide, 6.225% (w/v) **PC**, 0.2% (w/v) APS, 0.2% (w/v) TEMED], and immediately transferred to the gelation chamber in a humidified 37°C incubator for gelation. For the gelling solution, 4HT was replaced with purified water to ensure swift gelation. In addition, the gelation was performed for 1 hour instead of 2 hours.

For the spatially controlled photodegradation experiment (**Fig. 1E**), the gelled and digested cell samples were incubated in a SiR-DNA (CY-SC007, Cytoskeleton, Inc) solution (1:300 dilution with 1x PBS) for 1 hour prior to expansion and imaging.

*c. Mouse brain slices*

Photodegradable ~40 μm mouse brain slices were prepared for the two-photon photodegradation and isotropy experiments (**Fig. 2B-2D, fig. S5**) using the basic protocol.

A photodegradable gel block containing a ~40 μm mouse brain slice was prepared for the sequential two-photon photochemical sectioning experiment (**Fig. 2E-2G**) using a modified basic protocol. For the gelation chamber spacer, glass slides of ~1.0-1.2 mm in thickness were used instead of the coverslips. Furthermore, after expanding the sample in purified water, the gel block was trimmed into a cube with a side length of ~4-5 mm. The trimmed gel block was then rotated to stand up vertically (with ~4-5 mm in height) and immobilized on a coverslip using the poly-L-lysine mounting method for the subsequent imaging and photodegradation (*48*).

*d. Mouse olfactory bulbs (OBs)*

Photodegradable mouse OBs were prepared using a modified basic protocol. After immunostaining, the OBs were incubated in the AcX solution (0.1 mg/mL in 1x PBS) for 2 days, which was replaced with a freshly prepared AcX solution every 12 hours. The OBs were then incubated in the monomer solution (as in the basic protocol) at 4°C for 1-2 days and in the gelling solution (as in the basic protocol) at 0°C for 60 min before gelled in a humidified 37°C incubator for 2 hours. For the gelation chamber spacer, double-stacked glass slides (~2.0-2.4 mm in thickness) were used. After gelation, the samples were digested with proK for 2-3 days in the digestion buffer, which was replaced with a freshly prepared digestion buffer every day. The digested samples were stored in 1x PBS at 4°C for the subsequent imaging and photodegradation.

For the non-photodegradable OB serial sectioning experiments (**fig. S1**), immunostained OBs were (A) gelled using the standard protein-retention expansion microcopy protocol and serially sectioned to ~250 μm slices in 1x PBS using a vibratome (VT1200S, Leica), or (B) serially sectioned to ~100 μm slices using the same vibratome and gelled using the standard protein-retention expansion microscopy protocol. For the non-photodegradable OB block-face sectioning experiment (**fig. S2, Movie S1**), immunostained OBs were gelled using the standard protein-retention expansion microcopy protocol and sectioned using the same vibratome.

*e. Human hippocampus slices*

Photodegradable human hippocampus slices were prepared using a modified basic protocol. Briefly, the stained brain slices were incubated in the AcX solution (0.1 mg/mL in 1x PBS) for 2 days, which was replaced with a freshly prepared AcX solution every 12 hours. The brain slices were then incubated in a modified monomer solution [1x PBS, 2 M NaCl, 8.625% (w/v) sodium acrylate, 2.5% (w/v) acrylamide, 7.470% (w/v) **PC**] at 4°C for 1 day and in the same gelling solution at 0°C for 60 min before gelled in a humidified 37°C incubator for 2 hours. For the gelation chamber spacer, a stack of three no. 1.5 glass coverslips were used. After gelation, the samples were digested with proK for 2-3 days in the digestion buffer, which was replaced with a freshly prepared digestion buffer every day. Finally, the digested samples were incubated in 4% (w/v) SDS in 1x PBS at 37°C for 6-7 hours and washed with 1x PBS five times over two days. The samples were stored in 1x PBS at 4°C for the subsequent imaging and photodegradation.

*f. Fluorescently-labeled, photodegradable blank gel*

Fluorescently-labeled, photodegradable blank gel (blank PC-gel) was prepared using a modified basic protocol. Briefly, the monomer solution (as in the basic protocol) was mixed and incubated with an AcX and an Alexa Fluor 488 amine (218C0, Lumiprobe) solution to final concentrations of 2.1 and 5 mg/mL, respectively, at room temperature for more than 1 hour with shaking. Stock solutions of APS and TEMED were added to the mixed monomer solution to final concentrations of 0.2% (w/v) and 0.2% (w/v), respectively, to yield the gelling solution. The gelling solution was then transferred to gelation chambers of various shapes and sizes and gelled in a humidified 37°C incubator for 2 hours. Finally, the synthesized gels were stored in 1x PBS at 4°C for the subsequent use.

#### 4. Instrumentation for Sample Imaging and Photodegradation

*a. Confocal microscope*

An inverted Yokogawa spinning-disk (CSU-W1) confocal system on a Nikon Eclipse Ti2-E microscope body was used to image and photodegrade both cell and tissue samples. Briefly, single-photon photodegradation was performed using a 405 nm solid state laser and a CFI Apo LambdaS LWD 40x (1.15 NA) water-immersion objective. Sample imaging was performed using 488 nm, 561 nm, and 640 nm solid state lasers, a CFI Apo LambdaS LWD 40x (1.15 NA) water-immersion objective, and a Hamamatsu ORCA-Fusion BT C15440 sCMOS camera. Sample imaging and photodegradation were controlled by NIS-Elements AR v5.30.04 (Nikon).

*b. Multi-photon microscope*

An upright Leica SP8 DIVE multi-photon microscope with a HC IRAPO L 25x (1.00 NA) W motCORR water-dipping objective (11507704, Leica; WD: 2.6 mm) was used to image and photodegrade the mouse brain slice samples. For single-photon confocal imaging, green (488 nm), red (552 nm), and far-red (638 nm) channels were used to excite the fluorophores, and the Hybrid Detector (HyD) was used to detect the fluorescence signals. For two-photon photodegradation, an adjustable Spectra Physics Mai Tai laser (690-1040 nm) was used to illuminate the sample at a wavelength of 740 nm.

*c. Lattice light-sheet microscope*

Sequential on-block volumetric lattice light-sheet imaging and light-sheet photochemical sectioning was performed using a lattice light-sheet microscope similar to the one used for expansion lattice light-sheet microscopy (*7*, *49*). Briefly, a 488 nm laser (500 mW, 2RU-VFL-P-500-488-B1R, MPB Communications Inc.), 560 nm laser (1000 mW, 2RU-VFL-P-1000-560-B1R,

MPB Communications Inc.), and a 642 nm (2000 mW, 2RU-VFL-P-2000-642-B1R, MPB Communications Inc.) laser were expanded to $1/e^2$ diameter of 2.0 mm, combined into one path, and passed onto an acousto-optic tunable filter (AOTF; AOTFnC-400.650-CPCh-TN, Quanta-Tech, AA Opto Electronic). The collimated beam was fanned out to uniformly expand in the $x_{optical}$ axis using a Powell lens (LOCP-8.9R20-2.0, Laserline Optics Canada). The $z_{optical}$ axis was expanded using a pair of 50- and 250-mm cylindrical lenses (25 mm diameter; ACY254-050, LJ1267RM-A, Thorlabs). The expanded beam illuminated a horizontal stripe on a grayscale spatial light modulator (SLM; AVR17-0105, Meadowlark Optics, AVR Optics). The light diffracted by the SLM was focused onto a mask containing user-selected annuli of numerous sizes (Thorlabs Imaging) to block unwanted DC and higher diffraction orders. The light passing through the mask was reflected off a pair of galvanometer mirrors (6SD11226 and 6SD11587, Cambridge Technology, Novanta Photonics), which were conjugated to the back pupil of the excitation objective (TL20X-MPL, Thorlabs) and used to scan along the $x_{optical}$ and $z_{optical}$ axes. This leads to a lattice light sheet of ~200 µm wide along the image $y$-axis. The fluorescence generated by the specimen was collected through the detection objective (20×, 1.00 NA, 1.8 mm working distance, 421452-9800-000, Zeiss), projected onto a pupil-conjugate deformable mirror (DM; DM69, ALPAO) that corrects system aberrations and simultaneously imaged onto two sCMOS cameras (ORCA Fusion with 2304 x 2304 pixels, Hamamatsu Photonics). Appropriate dichroic (T600dcrb, Chroma) and emission filters (FF03-525/50-25 and FF01-538/685, Semrock) were used to separate fluorescent signals into the two cameras. An EM drive (MLS-3252 Electromagnetic Direct-Drive, SmarAct) was used to ensure a smooth and continuous movement of the sample holder along the x direction for the "sample scan" imaging acquisition. The y and z movements were respectively controlled by two linear stages (SLS-5252, SmarAct).

For light-sheet photochemical sectioning, a 405 nm light-sheet was introduced to the imaging region horizontally to the sample coverslip via a separate optical path. The 405 nm laser beam (100 mW, iBeam-Smart-405-S-BZ-1, Toptica) was expanded to a $1/e^2$ diameter of 2.0 mm, laterally fanned out to uniformly expand using a Powell lens (LOCP-8.9R20-2.0, Laserline Optics Canada), and axially compressed using a 250 mm cylindrical lens. This resulted in the formation of a light sheet that had a thickness ranging from ~70 to 100 µm over an area of ~18 x 18 mm, which was sufficient to cover the region of the sample that needed to be photodegraded while maintaining a desirable thickness for precise photochemical sectioning.

#### 5. Sample Imaging and Photodegradation

Unless otherwise noted, all the dimensions are provided at post-expansion scale in this section.

##### *a. Bulk photodegradation*

Photodegradable blank gels were prepared and expanded in purified water using the basic gelation protocol (without the digestion step) with a round gelation chamber. The expanded gels were then transferred to a UV chamber (365 nm, 60 W, PC-60-DJ, Phrozen Tech Co. LTD) for bulk photodegradation.

##### *b. Confocal imaging and single-photon photodegradation*

Photodegradable HEK 293 cell samples were expanded in purified water and immobilized onto a 6-well glass bottom well plate using the poly-L-lysine mounting method (*48*). The expanded samples were first imaged using the 488 nm, 561 nm, and 640 nm channels with a Z-step of 0.4-1 μm across a 2-by-2 or 3-by-3 tiled field-of-view (FOV, 15% overlap) on the confocal microscope (Yokogawa CSU-X1, Nikon). For photodegradation, the 405 nm channel was used to illuminate the central FOV for ~6-8 min (see **Table S1**). After illumination, the samples were imaged again using the 488 nm, 561 nm, and 640 nm channels with a Z-step of 0.4-1 μm across the same 2-by-2 or 3-by-3 tiled FOV.

For non-photodegradable controls, expanded non-photodegradable HEK 293 cell samples were subject to the same 405 nm illumination for >20 min and imaged across a 3-by-3 tiled FOV before and after the illumination.

To study the relationship between photodegradation and **PC** concentration, photodegradable HEK 293 cell samples were prepared with different **PC** concentrations [1.992% (w/v), 2.490%

(w/v), 3.735% (w/v), 4.358% (w/v), 4.980% (w/v), and 6.225% (w/v)] and subject to the same 405 nm illumination.

*c. Confocal imaging and two-photon photodegradation*

For the two-photon photodegradation experiment (**Fig. 2B-2D**), photodegradable mouse brain slices were expanded in purified water and immobilized on a no.1.5 coverslip glued to a 100 mm petri dish using the poly-L-lysine mounting method (*48*). The petri-dish containing the samples was then mounted to the multi-photon microscope (SP8 DIVE, Leica). The sample was then imaged using the microscope's single-photon confocal mode with green (488 nm), red (552 nm), and/or far-red (638 nm) channels at a Z-step size of 2 μm and a zoom factor of 1. Next, the middle volume (~60 μm thick) of the sample was photodegraded using the multi-photon mode at a wavelength of 740 nm at a Z-step size of 1 μm, a laser intensity of 30%, and a zoom factor of 2 for ~11 min. The samples were imaged again in the single-photon confocal mode using the same imaging conditions.

For the sequential two-photon photochemical sectioning experiment (**Fig. 2E-2G**), the photodegradable gel block containing the mouse brain slice was expanded, vertically rotated, and immobilized on a no. 1.5 coverslip glued to a 100 mm petri dish using the poly-L-lysine mounting method. Single-photon confocal imaging and two-photon photodegradation were then performed iteratively. Briefly, part of the top ~1.5 mm of the gel block containing the tissue was imaged in the single-photon confocal mode with XY tiling (two FOVs, 10% overlap) at a Z-step size of 5 μm and a zoom factor of 0.75. Next, the top ~1.0 mm of the gel block was photodegraded in the multi-photon mode at a wavelength of 740 nm with XY tiling (42 FOVs, 10% overlap), a Z-step size of 5 μm, a laser intensity of 100%, and a zoom factor of 0.75. After the two-photon photodegradation, no fluorescence signals were observed in the top ~1.0 mm of the gel block. Next, the plane lying ~1.0 mm below the initial gel surface was defined as the new surface of the gel, and another ~1.5 mm volume of the sample was imaged in the single-photon confocal mode with XY tiling (two FOVs, 10% overlap), a Z-step size of 5 μm, and a zoom factor of 0.75. The same process of two-photon photodegradation, redefinition of the gel surface, and single-photon confocal imaging (with the last imaging step covering a ~1.8 mm thick volume) was repeated once more until the total imaging depth from the initial gel surface reached ~3.8 mm.

*d. Light-sheet imaging and light-sheet photodegradation*

Photodegradable mouse OB and human hippocampus samples were expanded in 1x PBS and purified water, respectively, trimmed, and mounted on a cleaned 25 mm coverslip using the superglue mounting method with minor modifications (*48*). The 25 mm coverslips were soaked in 1M KOH in water for 30 min, rinsed with purified water three times, stored in 30% (v/v) ethanol in water, and air-dried before the sample mounting.

A modified superglue mounting method was used to immobilize the photodegradable mouse OB and human hippocampus samples to the 25 mm coverslip as it provided stronger and long-lasting adhesion. Briefly, a thin layer of superglue was applied to a small area of the cleaned 25 mm coverslip. Excessive liquid around the expanded sample was wicked away before the samples were placed on the applied superglue. After the gel was glued to the coverslip, it was submerged under 1x PBS or purified water for several hours for further curation of the superglue and for soaking away the impurities from the superglue. After curing, an opaque interface was formed between the gel and the superglue.

A modified poly-L-lysine mounting method was used to immobilize other samples (e.g., ~100-300 μm thick) to the 25 mm coverslip (*48*). Briefly, a few droplets of 0.1% (w/v) poly-L-lysine aqueous solution were applied to the top surface of the cleaned 25 mm coverslip for 20 min. The poly-L-lysine modified surface was rinsed with purified water three times and air-dried for 1 hour in a clean environment. Excessive liquid around the expanded samples was wicked away before the samples were placed on the poly-L-lysine modified surface of a 25 mm coverslip. After 20-30 s, a few droplets of 1x PBS or purified water were added to keep the sample hydrated.

Following sample immobilization onto the 25 mm coverslip, a 100 μL volume of 1:1000 diluted fluorescent bead solution [amine-terminated FluoSpheres, 0.2 μm, green fluorescent (488/515), Thermo Fisher] was applied to the sample surface for lattice light-sheet autofocusing. The coverslip was subsequently secured to the lattice light-sheet microscope sample holder using either superglue or metal clips. Finally, the sample holder was transferred to a sample chamber containing either 1x PBS or purified water.

Lattice light-sheet imaging of all samples was performed in the sample scan mode where the sample was translated continuously in the plane of the coverslip to allow for increased field of view and long-range fast scanning. To cover a large sample volume, we specified a rectangular parallelepiped as the hard limits for the tiled volume. The microscope software covered the volume with a 3D matrix of rectangular tiles with a desired tile overlap of 10 µm in sample Y axis and 3 µm in sample Z axis.

Before imaging, autofocus was performed on a 0.2 µm diameter fluorescent bead located on the sample surface. During imaging, autofocus was performed on a puncta fluorophore in the sample every ~8 hours to account for small system drifts. During each autofocus measurement, the bead or the puncta was precisely located using a normal imaging volume sweep. The light-sheet was statically held at the bead, while the sample stage was swept along the axis of the detection objective. The fluorescence intensity as function of a piezo position was fitted with a Gaussian curve and the peak center gave the correct piezo offset to use.

The lattice light-sheet applied in this study was a dithered HexRect lattice with a center numerical aperture of 0.25 and a Gaussian bounding factor of 0.08 (*49*). This resulted in an effective field-of-view of ~200 µm in the image y direction and ~70 µm in the image x direction. The microscope resolution for signals emitted at a wavelength of 500 nm was ~250 x 250 x 520 nm.

When possible, an additional sample mounting procedure was implemented to increase the imaging depth per cycle and minimize the deformation caused by scattered photodegradation light to the unimaged volume. Upon mounting the sample-laden coverslip onto the sample holder, the narrow axis of the sample was aligned parallel to the scan direction. This alignment situates the sample between the two objectives, thereby permitting a larger imaging depth as opposed to positioning the sample with its long axis parallel to the scanning direction.

In the case of mouse OB samples, which had an expansion factor of ~2 and a lateral scan range of ~6 mm, it was possible to reach an imaging depth of ~800 µm without the gel contacting the objectives. In practice, we limited the imaging depth to ~600 µm from the gel block-face. After imaging every subvolume, light-sheet photochemical sectioning with the 405 nm laser was performed till ~400 μm above the bottom plane of the imaged subvolume. We established this ~400 μm distance to minimize the impact of the scattered photodegradation light on the underlying sample while maintaining a reasonable thickness for each subvolume. The light-sheet illumination was applied for ~2-3 hours to complete the photodegradation. Subsequent imaging of the next sample subvolume commenced from ~150-200 μm above the bottom of the previously imaged one, ensuring sufficient overlap between the imaged subvolumes, and continued until the imaging reached a total thickness of ~600 μm. This overlap served as a reference for our computational stitching pipeline. In summary, each imaging-photochemical sectioning cycle encompassed volume with a total thickness of ~600 μm, including a ~150-200 μm thick overlap with the preceding subvolume and ~400-450 μm from the current subvolume. Additional imaging details can be found in **Table S1**.

The human hippocampus sample was imaged in purified water with an expansion factor of ~4.4 using similar experimental settings as the mouse OB samples. Briefly, each imaged subvolume extended across ~250 x 230 x 300 μm, including a ~100 μm overlap with the preceding one. A total of 8 subvolumes were imaged, resulting in a total volume of ~250 x 230 x 1630 μm (~57 x 52 x 370 μm at pre-expansion scale). Between each round of imaging, the same light-sheet illumination with the 405 nm laser was applied for ~20 minutes to complete the photodegradation. Additional imaging details can be found in **Table S1**.

#### 6. Image Preprocessing

##### a. *Flat-field correction, deconvolution, deskewing/rotation, and stitching*

For a subvolume imaged after each round of photochemical sectioning and imaging (including the first subvolume imaged before the photochemical sectioning), image tiles within the subvolume were initially stitched together into a subvolume using PetaKit5D software (*28*). During stitching, flat-field correction was applied to individual tiles as part of the preprocessing step. The flat-field and background images were estimated from these tiles. For each subvolume, the background image was estimated by averaging the blank frames from all tiles within the subvolume. The flat-field image was then estimated using the xy maximum intensity projections (MIPs) from all tiles along with the estimated background image through the BaSiC software (*50*).

After stitching, masks for the xy, xz, yz MIPs were generated from the stitched subvolume MIPs using user-defined thresholds to delineate object boundaries. Then, each subvolume was deconvolved with experimentally measured point-spread-functions (PSFs), followed by deskewing/rotation. Both processes used the large-scale processing strategies in PetaKit5D, employing the MIP masks to skip empty regions. The deskewing/rotation results were output with a Nyquist voxel size of 127. 5 x 127.5 x 168.5 nm for the mouse OB datasets.

After deskewing/rotation, the intensity profiles of the subvolumes were adjusted to correct uneven intensities resulting from the photobleaching and antibody gradient effects both within and across the subvolumes. To adjust the intensity profiles within each subvolume, the MIPs were used to estimate the intensity profiles along each axis. The xz MIP was initially smoothed using a Gaussian filter with a sigma of 2. Subsequently, the median value of all nonzero elements was calculated. The median values along the z-axis were calculated and normalized by dividing them by the overall median value. These normalized median values were then used for linear fitting. Correction factors along the z-axis were derived from this linear fitting model and capped to user-defined values (i.e., [0.5, 2] or [0.4, 2.5]) to avoid excessively extreme corrections. The correction factors along the x-axis were calculated using the median values along the x-axis. Similarly, the correction factors along the y-axis were determined using the yz MIP. Furthermore, an intensity correction factor for each subvolume was calculated by dividing a constant value (i.e., 5000) by the median intensity of the xy MIP. Intensity correction was applied using both axis-specific correction factors and an overall correction factor across subvolumes. These factors were multiplied and then limited to user-defined ranges (e.g., [0.4, 2.5] or [⅓, 3]). The intensity correction was performed using batch processing with a batch size of [1024, 1024, 1024], applying the correction factors to the corresponding batches.

After intensity correction, the scales on the x and y axes of the subvolumes were adjusted to account for the sample's varying expansion during acquisition or gel degradation. The xz and yz MIPs were used to determine the optimal resampling factors between adjacent subvolumes.

Ultimately, these pairwise resampling factors were unified across all subvolumes. The resampling tool in PetaKit5D was used to rescale the subvolumes with the unified resampling factors.

After resizing each subvolume along the x and y axes, the next step was to stitch them together. Initially, subvolume coordinates were estimated using the overlapped volumes identified from the MIPs. Rigid 3D stitching was performed using the large-scale strategy described in PetaKit5D with cross-correlation registration and feather blending. Here, bounding boxes to define dominant regions were applied during feather blending to assign higher weights to the overlapped volumes from the top subvolumes (which were imaged earlier) to better preserve the image quality, as the overlapped volumes from the bottom subvolumes were subject to an increased photobleaching effect. To stitch the WT and MT mouse OB datasets, the registration information from the full-resolution and 5 x 5 x 5 downsampled data were used for the stitching, respectively. We found the latter provided sufficient performance for registration.

The processing of the non-photodegradable serially sectioned mouse OB samples followed a similar procedure. The subvolumes, whose axes were flipped randomly during sample preparation and mounting during acquisition, were corrected to ensure consistent orientation across all the subvolumes. The cross-correlation registration was applied to refine the positions of all subvolumes during stitching, even in the absence of overlapping volumes between them.

The human samples were processed similarly to the mouse OB datasets, excluding the intensity correction and xy resampling. Steps included flat-field correction, stitching of each subvolume, deconvolution, deskewing/rotation, stitching across the subvolumes, and exported at a voxel size of 98.0 x 98.0 x 90.0 nm. All three channels were used for stitching within each subvolume, but only the axon (NF-200) and myelin sheath (MBP) channels were for stitching across the subvolumes.

The stitching of the sequential two-photon photochemical sectioning dataset (**Fig. 2E-2G**) followed a similar procedure. The tile coordinates were first estimated by manually inspecting intensity profiles across adjacent tiles and saved in an image list CSV file. Then the tiles were stitched together using the stitcher in PetaKit5D with the image list CSV file.

*b. Filtering non-specific antibody signals*

Non-specific antibody signals were filtered by removing small objects using variable background segmentation. There were two main steps: estimating local background and segmentation. In the first step, the local background within a volume of 128 x 128 x 128 voxels in size was estimated using batch processing with this batch size and an additional 64 voxel size border buffer on both sides. For each batch, the image was first smoothed with a 3D Gaussian filter with a user-defined sigma value (4 for the datasets). After smoothing the image, the xy MIP was computed. The candidate threshold value (T) was calculated based on the non-zero elements of the smoothed 3D data that fell below the 99.9th percentile in the xy MIP. Next, the xy MIP was used to determine another candidate threshold value (T_mask) by masking objects in the MIP. The mask for objects was generated by thresholding the background image, computed using a Gaussian filter with sigma 80 applied to the MIP. If the masked area covers more than 90% of the image, the multiplier applied to the background image was increased. Next, the background mask was generated by combining the object mask's complement after a 5-pixel erosion, with the MIP thresholder using user-defined minimum and maximum thresholds. This was followed by image opening and an additional 5-pixel erosion. After obtaining the background mask, the threshold T_mask was determined using the mode of voxels within the mask in the MIP. If this threshold is lower than the 25th percentile, the median value is used instead. T_mask was then capped to [T_base, T_max]. The background value for the batch was defined as 90% of the mask threshold T_mask, plus 10% of the maximum value between the Otsu-based threshold (T) and T_max. After processing all batches, background values were collected at their corresponding locations within the 128 x 128 x 128 downsampled data.

In the second step, the processing was split into small batches of 1024 x 1024 x 1024 voxels, each with an additional border buffer of 100 voxels on both sides of all axes. For each batch, the local background region was loaded and up-sampled to match the target batch region using linear interpolation. Then the batch region was Gaussian smoothed with a sigma of 4 and binarized using the up-sampled background multiplied with a user-defined factor (i.e. 1.8). Objects in the binarized mask smaller than a user-defined volume threshold (e.g., 5000 for myelin and 2000 for axon channels) were discarded. The cleaned mask was applied to the raw region to eliminate small objects and backgrounds. After cropping out the border buffers, the processed data was saved to disk. The entire process was parallelized using the generic computing framework in PetaKit5D.

*c. Dataset export (for visualization)*

To balance fluorescence intensities across the datasets, adjustments were made using manually annotated masks defining five anatomical layers of mouse olfactory bulbs [generated with ITK-SNAP (ver. 4.0.1)]: olfactory never layer (ONL, including the surrounding tissue), glomerular layer (GL), external plexiform layer (EPL) and mitral cell layer (MCL), internal plexiform layer (IPL), and granule cell layer (GCL). For each layer, the upper bound for normalization was determined by the 99.99th percentile of non-zero voxel intensities in the 20 x 20 x 20 downsampled data, with 0 as the lower bound. The data at full resolution was normalized using these upper and lower bounds, then rescaled to a range of 0 to 65535 for each layer. These normalization processes were performed using batch processing with the generic computing framework in PetaKit5D.

For Imaris visualization, the 5 x 5 x 5 downsampled data were normalized as described above. The normalized data was converted to an Imaris file using the Imaris File converter in PetaKit5D.

#### 7. Image Analysis

*a. Segmentation and skeletonization of axons and myelin sheaths*

To segment the axons and myelin sheaths, a procedure akin to filtering non-specific antibody signals was applied to the final stitched data (before removing the non-specific signals). Specifically, we used a threshold multiplier of 2.0 for local background and applied Gaussian smoothing with a sigma of 2.5.

After segmenting the data, skeletonization was performed using batch processing with a size of 1024 x 1024 x 1024 voxels and borders of 100 x 100 x 100 voxels on the segmentation. The MATLAB function bwskel was applied to each batch, setting a minimum branch threshold of 50 voxels to eliminate small branches. Following this, the skeletonized blobs were pruned by removing clusters with 5 or more neighboring voxels, along with their associated neighboring voxels.

We note that features of an extremely dim and punctuated nature can be removed by this step. For example, some of the dim and punctuated axon segments were removed and consequently excluded from the axon length and myelination analysis, which could have led to an underestimation of the total axon length and associated parameters. Improving the immunostaining quality and/or keeping the expansion factor at a reasonably low range (to maintain a high labeling density) can potentially mitigate this problem.

*b. Axon and myelination analysis*

The axon skeletonization and myelin mask were used to access myelination of axons. An axon voxel was considered myelinated if it intersected with the myelin mask or fell within a user-defined threshold (e.g., 1 μm) of the nearest myelin voxel. Batch processing with a batch size of 1024 x 1024 x 1024 voxels was used for the analysis. For each batch, the images were first up-sampled to isotropic voxel size (i.e., 127.5 x 127.5 x 127.5 nm) via nearest neighbor interpolation. Next, myelinated voxels were identified, and total and myelinated axon lengths were computed for each layer. After processing all batches, the total and myelinated axon lengths were summarized over the results of all batches for each layer. The myelination ratios were calculated by dividing the myelinated axon lengths by the corresponding total axon lengths.

To determine the axonized myelin, that is, the myelin with axons inside, a similar process was conducted using the myelin skeletonization and axon mask.

As described, we excluded ONL from our OB analysis and quantification because this layer contained additional tissue from the OB surface or the surrounding structures and had elevated level of immunostaining background. Hence, the masked region did not provide an accurate representation of neuronal structures within the layer.

*c. Multi-region of interest (ROI) analysis*

50 cropped ROIs of 1000 x 1000 x 1000 voxels (~66 x 66 x 87 μm at pre-expansion scale) from each layer were generated from an evenly spaced grid for the centers across the entire volume of the dataset. The number of grid points in each dimension was approximately proportional to the dimensions of the volume. Briefly, an initial set of grid point numbers was used to generate the grid coordinates for the centers throughout the entire volume, with the largest possible grid distances to cover the entire volume and the grid centered. Then, each ROI centered at a grid coordinate was checked to see if it was fully contained within the layer, as defined by the manually annotated masks. For the IPL layer, the constraint was relaxed to include an ROI with at least 75% of the volume within the layer, as this layer was thin. The valid ROIs following the above criteria were counted. If their number was between 50 and 55, the process stopped and the center coordinates for the valid ones were used to crop the ROIs. Otherwise, the grid was adjusted by changing the number of points to get the valid regions within the 50 to 55 range. This process was repeated for all the layers.

After identifying the centers of valid ROIs for all layers, the ROIs were cropped, and the XY MIPs of each ROI were generated for visualization at full resolution. The same method used for the analysis of the whole dataset was applied to the analysis of axons and myelin sheaths for each ROI.

*d. Tracing*

Semi-automatic tracing of axons was performed using Imaris Filament Tracer (ver. 9.3.1, Oxford Instruments) with the "AutoPath" method.

**8. Visualization**

Unless otherwise noted, all the 3D rendered datasets were flat-field corrected, deconvolved, stitched, filtered to remove non-specific antibody signals, and gamma adjusted for visualization.

The mouse OB datasets were visualized and 3D rendered at 5 x 5 x 5 downsampled resolution using Imaris (ver. 10.0.0, Oxford Instruments). The human hippocampus dataset was visualized and 3D rendered at full resolution using Imaris (ver. 9.3.1, Oxford Instruments). The 10 traced axons were displayed using the “Cone” style with a scale of 1, and the starting points and terminal points were indicated by blue and green balls, respectively.

Unless other noted, all the 2D cross-sectional and maximum intensity projection (MIP) views of the imaged volumes were generated using ImageJ distribution Fiji (ver. 1.54f).

#### Supplementary Figures

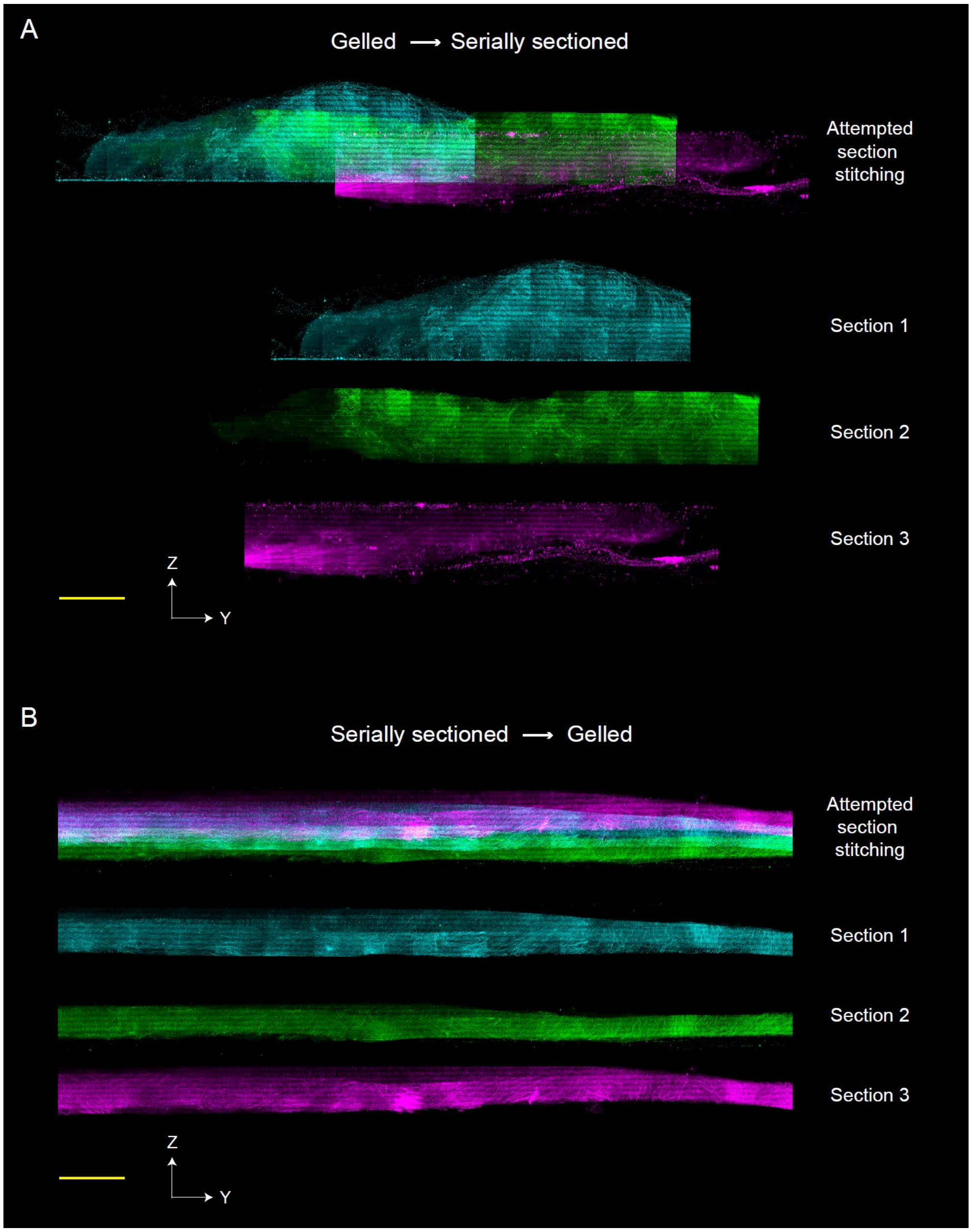


**Fig. S1. Fluorescence imaging and stitching of (A) hydrogel-embedded and serially vibratome-sectioned, and (B) serially vibratome-sectioned and hydrogel-embedded tissue (mouse olfactory bulb) samples.** For **A**, the tissue was fluorescently labeled for axons (NF-200, green) and myelin sheaths (MBP, red), embedded and expanded with polyacrylamide/sodium polyacrylate hydrogel with the bis-acrylamide crosslinker (Bis-gel), and serially sectioned using a vibratome (VT1200S, Leica; ~250 µm thick) in 1x PBS. For **B**, after the same fluorescence labeling, the tissue was serially sectioned using the same vibratome (~100 µm thick), and embedded and expanded with the Bis-gel in 1x PBS. All the serial sections were imaged in 1x PBS using a lattice light-sheet microscope and visualized in YZ maximum intensity projection (MIP) view. Scale bars: 50 µm (post-expansion scale).

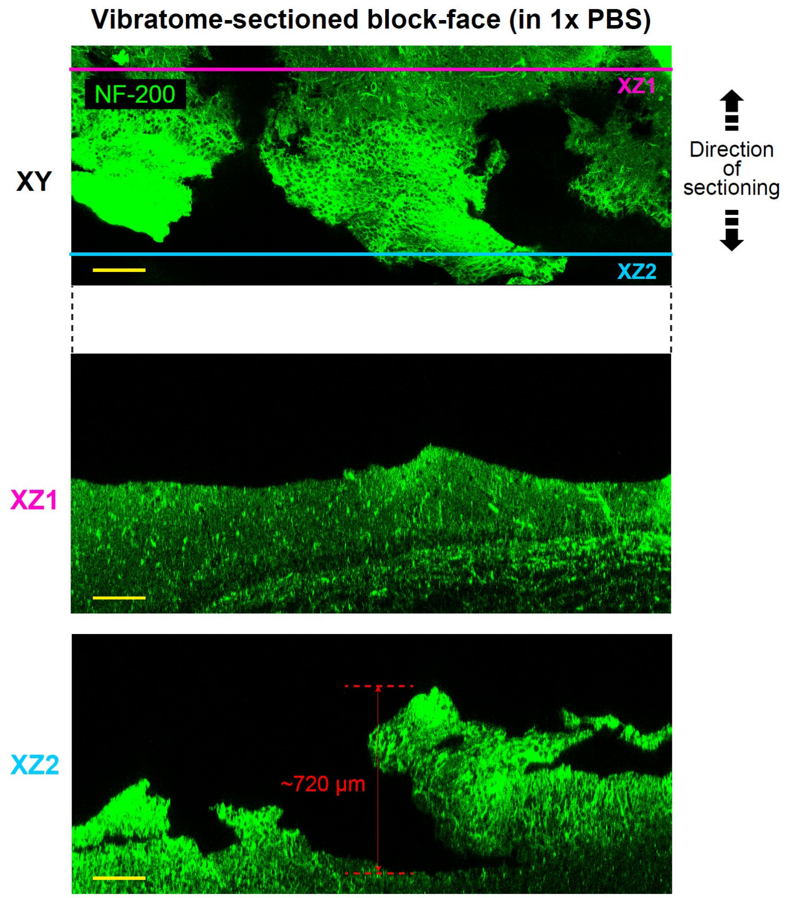


**Fig. S2. XY (top) and two XZ (bottom, XZ1 and XZ2) views of a vibratome-sectioned block-face of a hydrogel-expanded tissue (mouse olfactory bulb) sample.** The tissue was fluorescently labeled for axons (NF-200, green), expanded with the Bis-gel, and sectioned using a vibratome (VT1200S, Leica; ~300 µm thick) in 1x PBS (~2-fold expansion). The block-face was imaged in 1x PBS using the single-photon confocal mode on an upright two-photon fluorescence microscope. Scale bars: 200 µm (post-expansion scale).

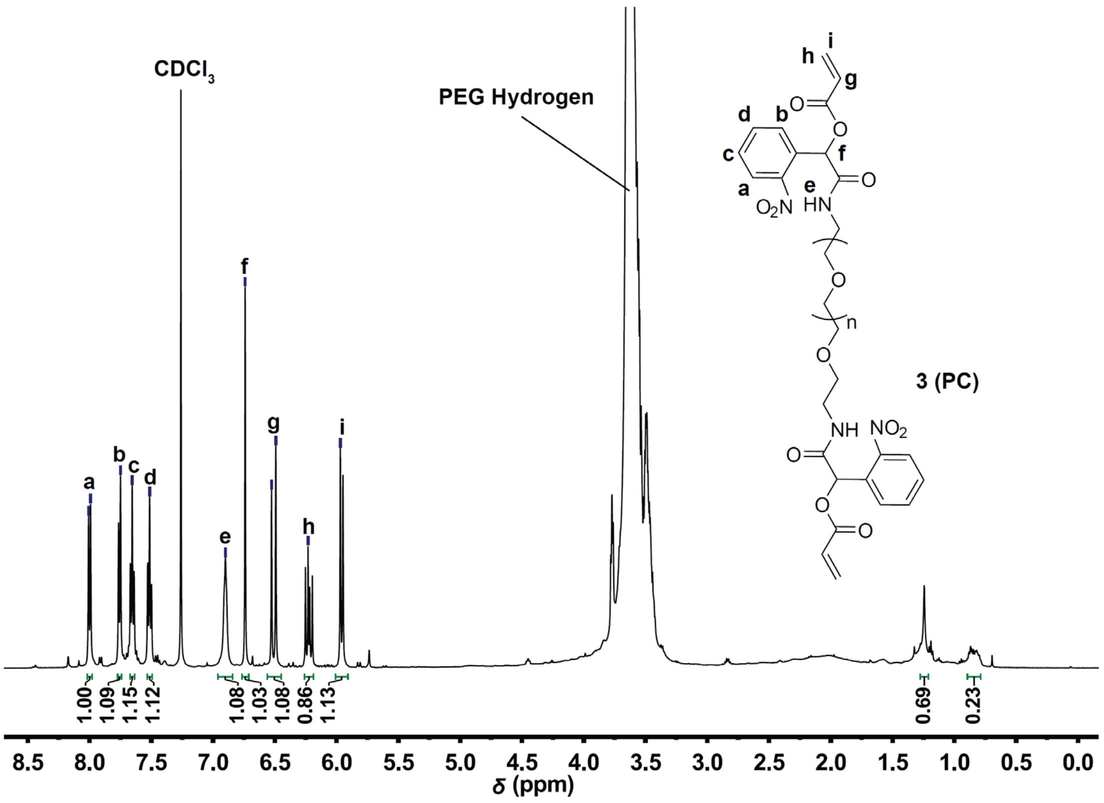


**Fig. S3.** $^{1}H$ **NMR spectra of compound 3 (PC)**.

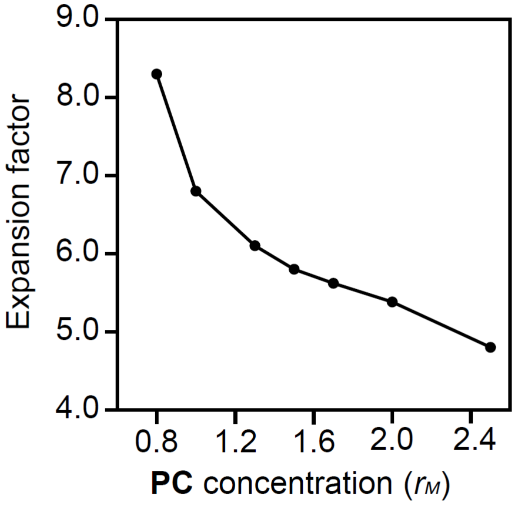


**Fig. S4. Expansion factor of photocleavable crosslinker (PC)-crosslinked polyacrylamide/sodium polyacrylate hydrogel (PC-gel) at different PC concentrations.** $r_M$, molar ratio of PC to bis-acrylamide, used in its non-photodegradable counterpart, bis-acrylamide-crosslinked polyacrylamide/sodium polyacrylate gel (Bis-gel).

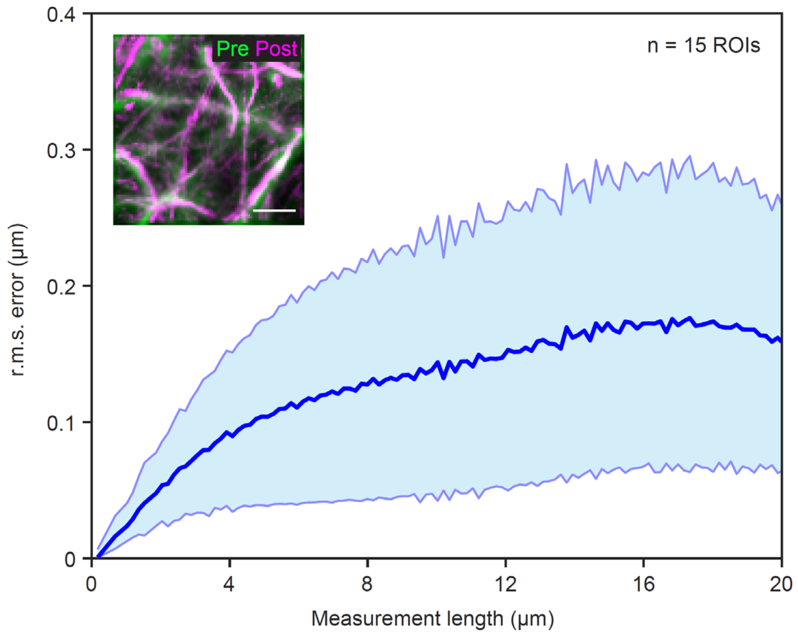


**Fig. S5. Expansion isotropy analysis.** Root-mean-square (r.m.s.) expansion error of ~40 µm thick mouse brain slices embedded and expanded with the PC gel [blue line, mean; shaded area, standard deviation; n = 15 regions of interest (ROIs) from three brain slices from one animal]. Inset: Non-rigidly registered and overlaid pre- (green) and post-expansion (magenta) images used for the r.m.s. error analysis. Scale bar, 3 µm (16.2 µm). Here and after, unless otherwise noted, scale bars are provided at pre-expansion scale (with the corresponding post-expansion size indicated in brackets).

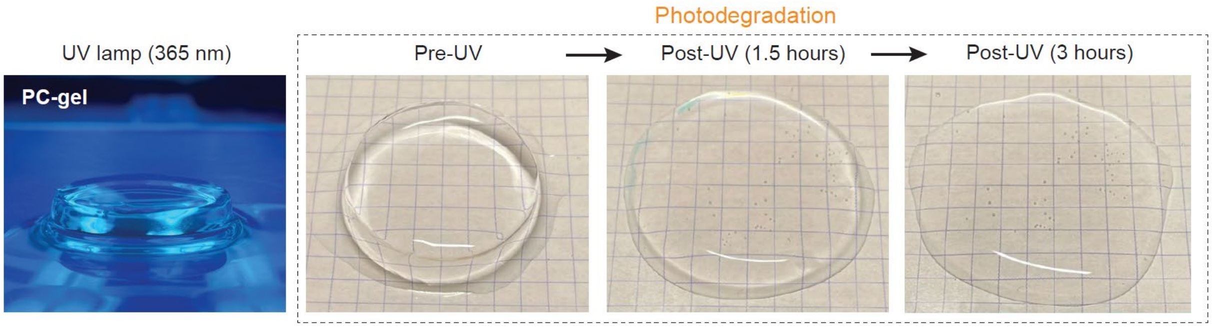


**Fig. S6. Bulk photodegradation of PC gel with UV lamp.** (Left) Image of PC-gel under UV lamp illumination (365 nm). (Right) Pre-UV, post-UV (1.5 hours), and post-UV (3 hours) images of the same PC-gel (expanded in purified water) under continuous UV illumination. Grid size: 5 mm.

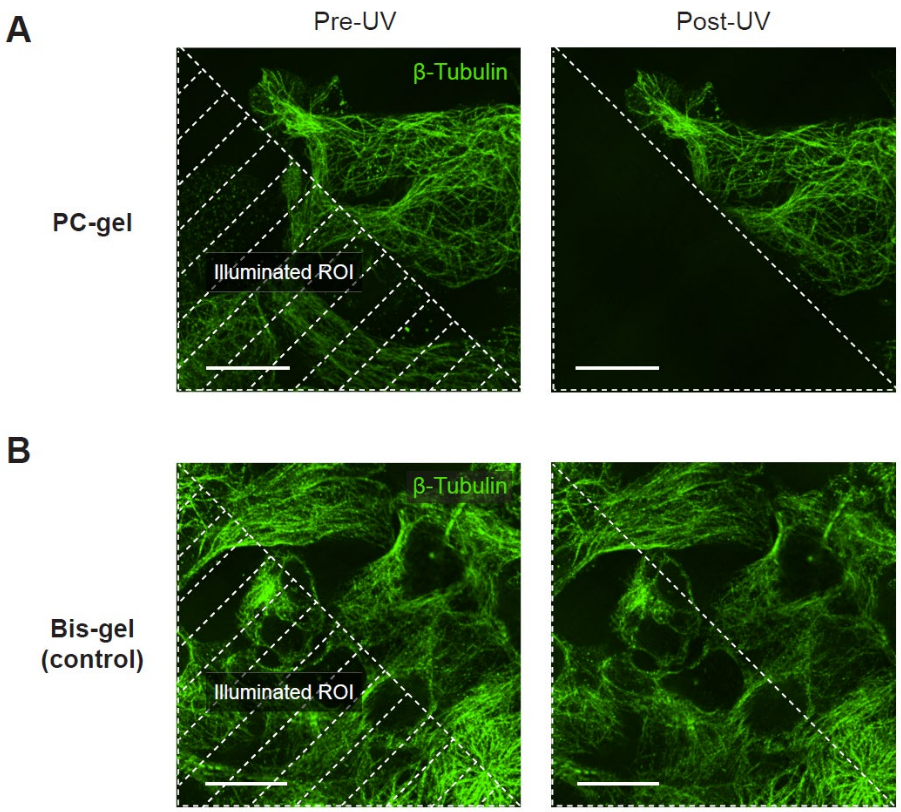


**Fig. S7. Spatial selective photodegradation of HEK cells labeled with green fluorescent dyes using a 405 nm UV laser.** (**A**) Pre- (left) and post-UV (right) images of HEK cells, fluorescently labeled for β-tubulin using AF488-conjugated antibodies and expanded with the PC-gel. The shaded area in the pre-UV image indicates the ROI for the 405 nm laser illumination. Scale bars: 10 µm (48 µm). (**B**) Pre- (left) and post-UV (right) images of HEK cells, fluorescently labeled for β-tubulin using AF488-conjugated antibodies and expanded with the Bis-gel (as control). The shaded area in the pre-UV image indicates the ROI for the 405 nm laser illumination. The 405 nm laser intensity was kept constant across **A** and **B**. Scale bars: 10 µm (45 µm).

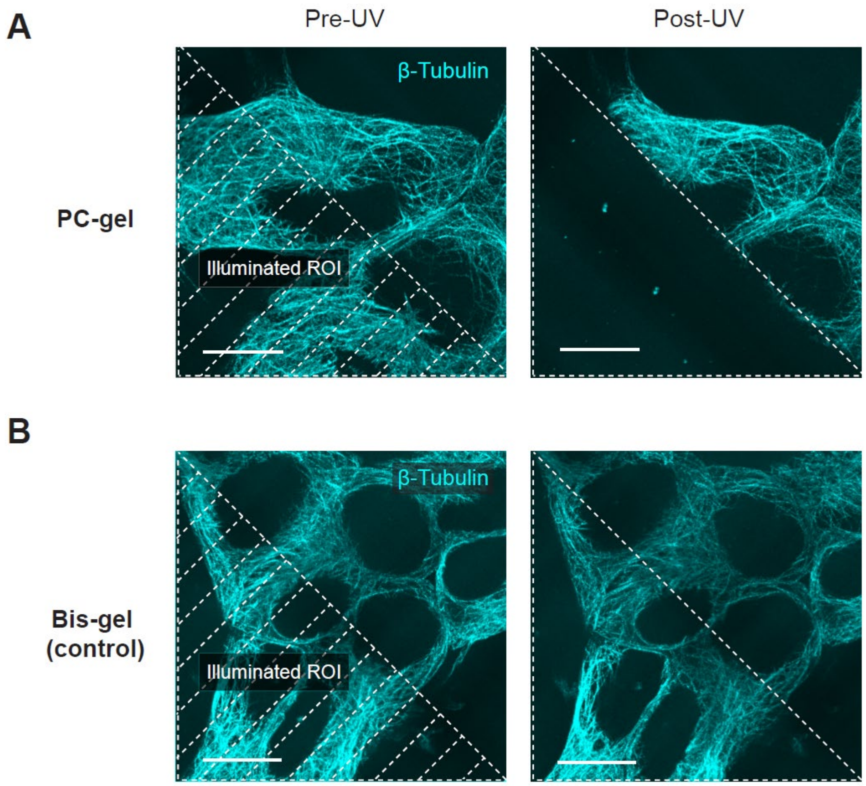


**Fig. S8. Spatial selective photodegradation of HEK cells labeled with far-red fluorescent dyes using a 405 nm UV laser.** (**A**) Pre- (left) and post-UV (right) images of HEK cells, fluorescently labeled for β-tubulin using ATTO 647N-conjugated antibodies and expanded with the PC-gel. The shaded area in the pre-UV image indicates the ROI for the 405 nm laser illumination. Scale bars: 10 µm (48 µm). (**B**) Pre- (left) and post-UV (right) images of HEK cells, fluorescently labeled for β-tubulin using ATTO 647N-conjugated antibodies and expanded with the Bis-gel (as control). The shaded area in the pre-UV image indicates the ROI for the 405 nm laser illumination. The 405 nm laser intensity was kept constant across **A** and **B**. Scale bars: 10 µm (45 µm).

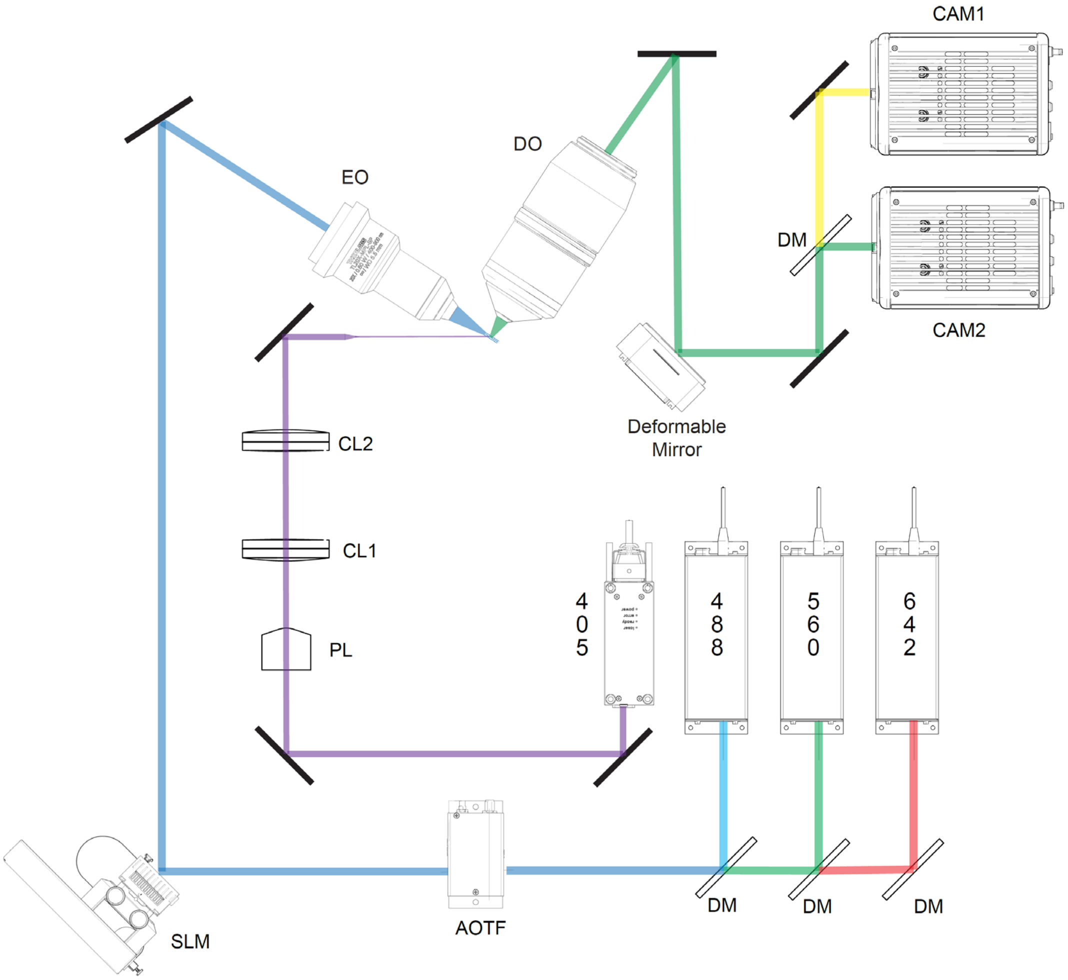


**Fig. S9. Simplified schematic showing the lattice light-sheet excitation, two-color detection, and light-sheet photochemical sectioning optical paths for light-sheet VIPS.** EO, excitation objective; DO, detection objective; CAM, camera; DM, dichroic mirror; CL, cylindrical lens; PL, Powell lens; SLM, spatial light modular; AOTF, acousto-optic tunable filter.

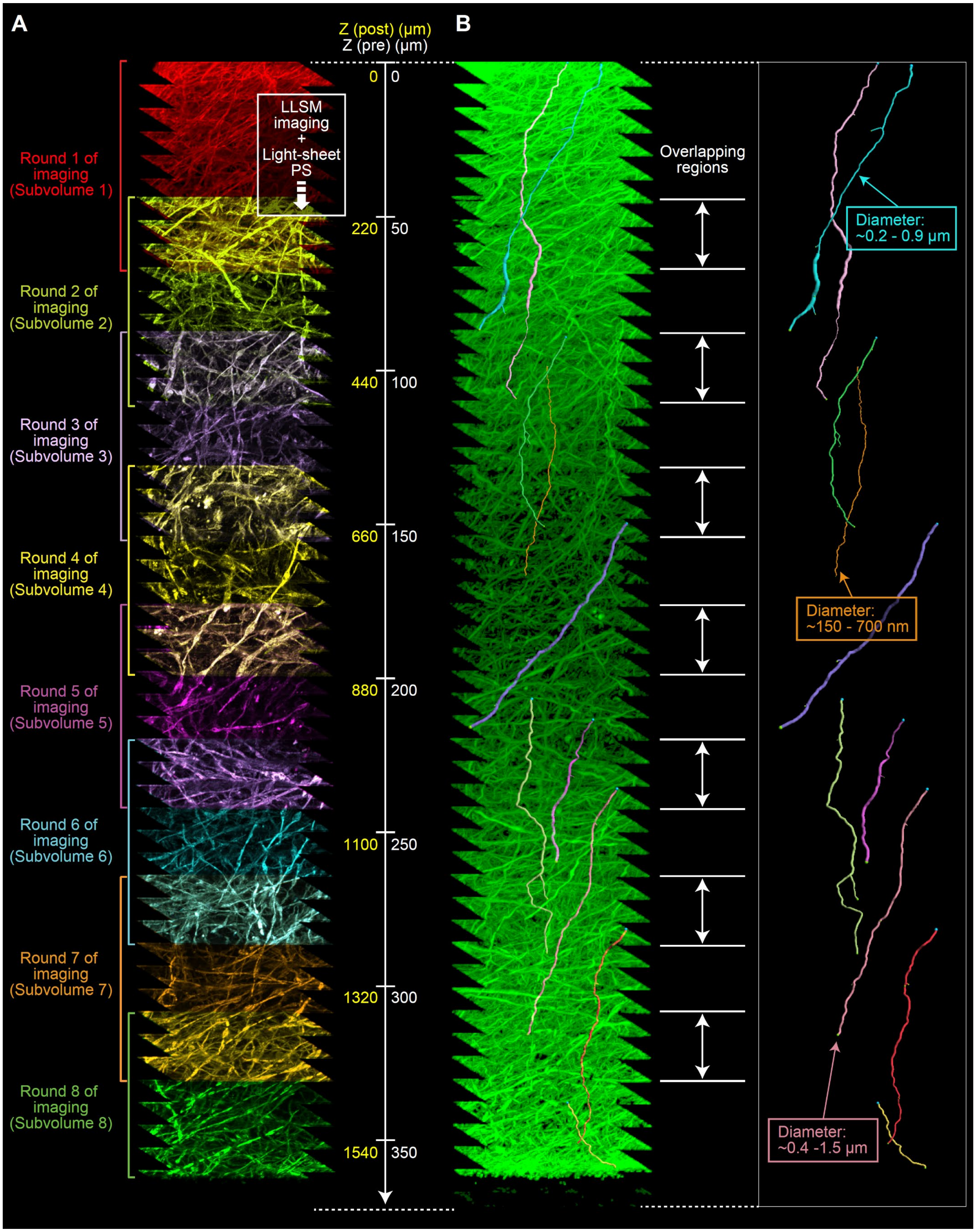
A
B
Z (post) (μm)
Z (pre) (μm)
Round 1 of imaging (Subvolume 1)
Round 2 of imaging (Subvolume 2)
Round 3 of imaging (Subvolume 3)
Round 4 of imaging (Subvolume 4)
Round 5 of imaging (Subvolume 5)
Round 6 of imaging (Subvolume 6)
Round 7 of imaging (Subvolume 7)
Round 8 of imaging (Subvolume 8)
LLSM imaging + Light-sheet PS
0 0
220 50
440 100
660 150
880 200
1100 250
1320 300
1540 350
Overlapping regions
Diameter: ~0.2 - 0.9 μm
Diameter: ~150 - 700 nm
Diameter: ~0.4 -1.5 μm

**Fig. S10. Nanoscale fluorescence imaging of human hippocampus with light-sheet VIPS.** (**A**) 3D rendered XZ slab view (~60 µm thick at post-expansion scale) of the stitched human hippocampus dataset, imaged with 8 rounds of sequential lattice light-sheet microscopy (LLSM) imaging and light-sheet photosectioning (with no sectioning after the last round of imaging). The axon (NF-200) channel of each imaged subvolume is distinctly colored for visualization purposes. The Z axis is shown at both post-expansion (yellow) and pre-expansion (white) scales. The imaged subvolumes were computationally stitched using the 3D rigid registration pipeline. (**B**) (Left) 3D rendered XZ view of the axon (NF-200) channel of the same human hippocampus dataset. Individual axons with diameters ranging from ~150 nm to 1.5 µm were traced from one to the other edge of the volume across multiple imaged subvolumes. Volumetric overlaps between the imaged subvolumes are indicated with the double-arrows. (Right) 3D rendered XZ view of the traced axons. The diameters of three individually traced axons are shown as an example.

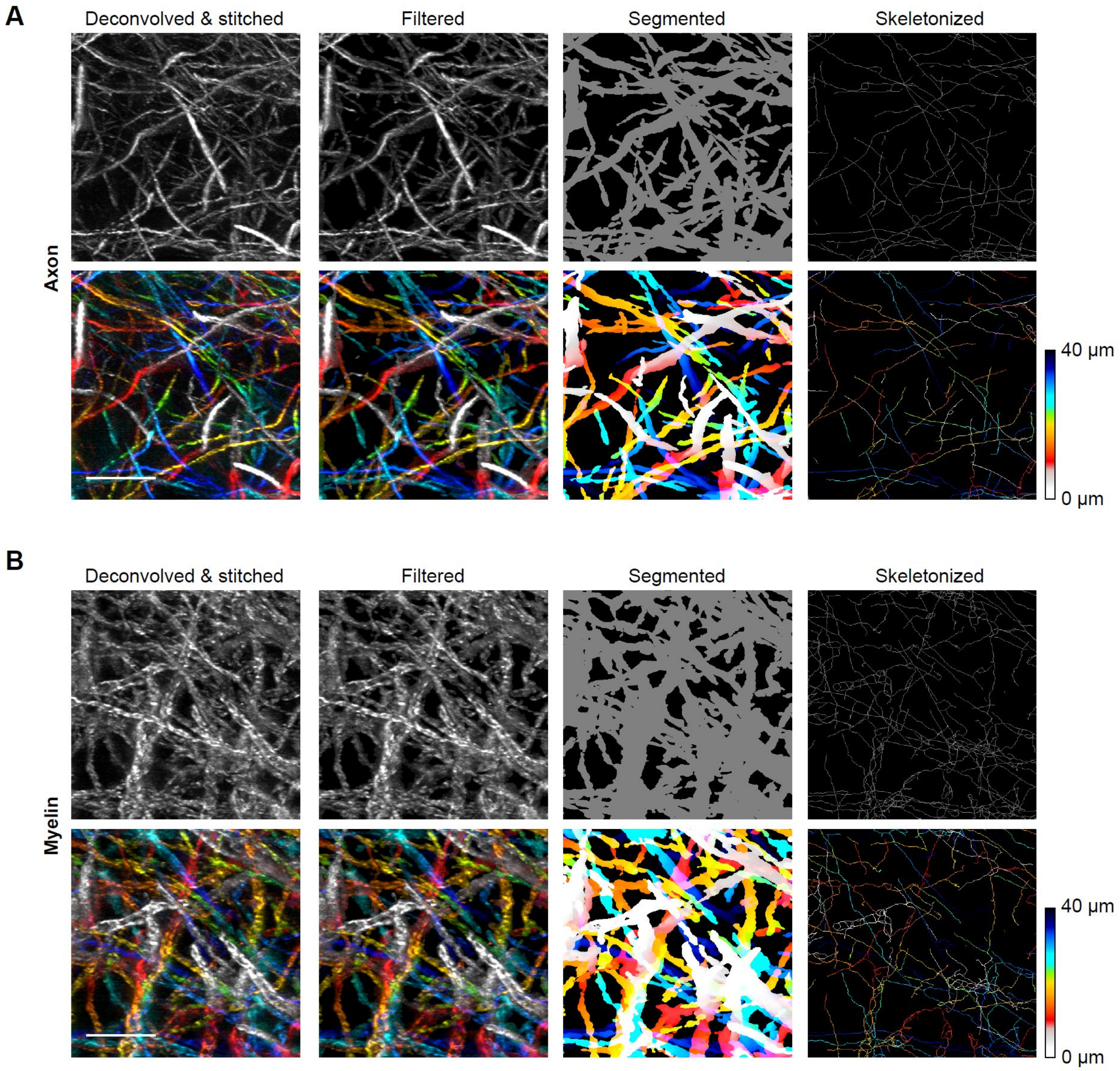


**Fig. S11. Segmentation and skeletonization of axons and myelin sheaths.** MIP view of the (**A**) axon (NF-200) and (**B**) myelin (MBP) channel of the same ROI of the wildtype (WT) mouse olfactory bulb (OB) dataset after deconvolution and stitching, filtering, segmentation, and skeletonization. The top and bottom row shows the same MIP image in greyscale and color-coded view (for Z), respectively. Scale bars, 10 (19.3) µm

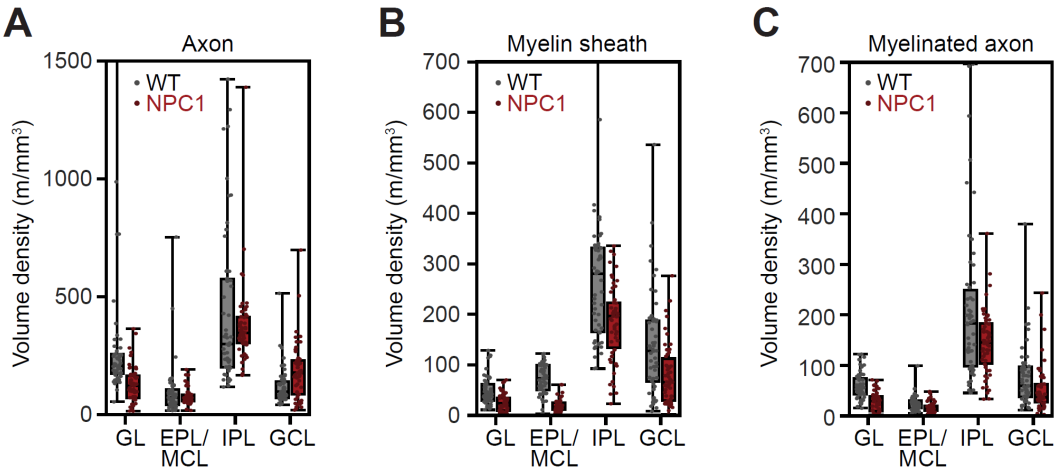


**Fig. S12. Volume density of axons, myelin sheaths, and myelinated axons across 50 ROIs.** Distribution of volume density of (**A**) axons, (**B**) myelin sheaths, and (**C**) myelinated axons across 50 ROIs (~66 by 66 by 87 µm at pre-expansion scale) randomly sampled across distinct anatomical layers of the WT and NPC1 mouse OB. Data are presented as box plots, where the ends of the whiskers represent the maximum and minimum values of the distribution, the dots represent individual values, and the upper, middle, lower line of the box represents the 75th percentile, 50th percentile (median), and the 25th percentile, respectively. ONL: olfactory nerve layer; GL: glomerular layer; EPL: external plexiform layer; MCL: mitral cell layer; IPL: internal plexiform layer; GCL: granule cell layer.

### Supplementary Tables

**Table S1.** Sample preparation, imaging, and image processing conditions.

| Sam ple # | Related Figure # | Sample Name | Sample Preparation | | | | Imaging and Photodegradation Conditions | | | | | | | Image Preprocessing | Image Analysis & Visualization |
|---|---|---|---|---|---|---|---|---|---|---|---|---|---|---|---|
| | | | **Genotype** | **Fluorescent proteins or target proteins/ epitopes for immunostaining** | **Antibodies** | **Expansion factor (gel-type) (medium)** | **Imaging/ Photode gradatio n mode** | **Effective voxel size (x, y, z) [post-expansion size in brackets] (nm)** | **Total imaged size (x, y, z) [post-expansio n size in brackets] (µm)** | **Imaging channels (exposure time; power)** | **Photodegradat ion channel (power)** | **Total imaging time (total photodegrad ation time) (hours)** | | **Flat-field, deconvolution, deskewing/rot ation, stitching, and additional corrections** | **Analysis pipelines and visualization parameters** |
| **1** | **Fig. 1E** | **Human cell (HEK cells)** | HEK | Beta-tubulin | Primary: rabbit polyclonal (ab6046, Abcam) Secondary: goat anti-rabbit Alexa Fluor 488 (A11008, Thermo Fisher) | 4.80 (PC-gel) (water) | Confocal/ Confocal | 34 x 34 x 83 [161 x 161 x 400] | NA | 488 (200 ms; 30%, 2.04 x $10^{-5}$ mW/µm$^2$) | 405 (100%, 5.17 x $10^{-5}$ mW/µm$^2$) | 0.38 for imaging (0.1 for photodegrada tion) | | NA | NA |
| | | | | DNA | SiR-DNA (CY-SC007, Cytoskeleton, Inc) | | | | | 640 (300 ms; 40%, 3.13 x $10^{-5}$ mW/µm$^2$) | | | | | |
| **2** | **Fig. S7** | **Human cell (HEK cells)** | HEK | Beta-tubulin | Primary: rabbit polyclonal (ab6046, Abcam) Secondary: goat anti-rabbit Alexa Fluor 488 (A11008, Thermo Fisher) | 4.80 (PC-gel) (water)<br>4.50 (Bis-gel) (water) | Confocal/ Confocal | 36 x 36 x 222 [161 x 161 x 1000] | NA | 488 (200 ms; 30%, 2.04 x $10^{-5}$ mW/µm$^2$) | 405 (100%, 5.17 x $10^{-5}$ mW/µm$^2$) | 0.22 for imaging the whole z-stack (0.1 for photodegrada tion; 0.33 for photobleachi ng of Bis-gel) | | NA | NA |
| **3** | **Fig. S8** | **Human cell (HEK cells)** | HEK | Beta-tubulin | Primary: rabbit polyclonal (ab6046, Abcam) Secondary: goat anti-rabbit ATTO 647N 40839-1ML-F, Millipore Sigma) | 4.80 (PC-gel) (water)<br>4.50 (Bis-gel) (water) | Confocal/ Confocal | 36 x 36 x 222 [161 x 161 x 1000] | NA | 640 (300 ms; 40%, 3.13 x $10^{-5}$ mW/µm$^2$) | 405 (100%, 5.17 x $10^{-5}$ mW/µm$^2$) | 0.15 for imaging the whole z-stack (0.1 for photodegrada tion; 0.33 for photobleachi ng of Bis-gel) | | NA | NA |
| **4** | **Fig. 2B-2D** | **Mouse brain slice** | C57BL/6 | Homer1 | Primary: rabbit polyclonal (160 003, Synaptic Systems) Secondary: goat anti-rabbit ATTO 647N 40839-1ML-F, Millipore Sigma) | 5.40 (PC-gel) (water)<br>(note: all imaging parameters reported in post-expansion size) | Confocal/ Two-photon | [867 x 867 x 2000] | [~440 x 440 x 135] | 638 (1.28 s/frame; NA) | 740 (30%, 1.10 x $10^{-3}$ mW/µm$^2$; zoom factor = 2) | 0.03 for imaging (0.2 for photodegrada tion) | | NA | NA |
| **5** | **Fig. 2E-2G** | **Mouse brain slice** | C57BL/6 | NF-200 | Primary: rabbit polyclonal (N4142-.2ML, Millipore Sigma) Secondary: goat anti-rabbit Alexa Fluor 488 (A11008, Thermo Fisher) | 5.40 (PC-gel) (water)<br>(note: all imaging parameters reported in post-expansion size) | Confocal/ Two-photon | [1156 x 1156 x 5000] | [~600 x 1260 x 3800] | 488 (1.28 s/frame; NA) | 740 (100%, 5.32 x $10^{-4}$ mW/µm$^2$; zoom factor = 0.75) | 2 for imaging (6 for photodegrada tion) | | Tile position estimation, stitching | NA |

| | | | | | | | | | | | | | | |
|---|---|---|---|---|---|---|---|---|---|---|---|---|---|---|
| | | | | MBP | Primary: chicken polyclonal (PA1-10008, Thermo Fisher) Secondary: goat anti-chicken Alexa Fluor 568 (A11041, Thermo Fisher) | | | | | 552 (1.28 s/frame; NA) | | | | |
| 6 | **Fig. 3 Fig. 4 Fig. S12 Movie S4** | **Mouse olfactory bulb (wild-type) “WT”** | BLAB/c (*Npc1*$^{+/+}$) | NF-200 | Primary: rabbit polyclonal (N4142-.2ML, Millipore Sigma) Secondary: goat anti-rabbit Alexa Fluor 488 (A11008, Thermo Fisher) | 1.93 (PC-gel) (1x PBS) | Lattice light-sheet/ Light-sheet | 56 x 56 x 135 [108 x 108 x 260] | ~3320 x 4870 x 2220 [~6410 x 9401 x 4290] | 488 (6 ms; 1.6 mW) | 405 (100 mW) | 142 for imaging (20 for photodegradation) | Flat-field correction, deconvolution, deskewing/rotation, intensity correction, xy-resampling, stitching, puncta removal Voxel size after deskewing/ rotation (nm): 66 x 66 x 87 [128 x 128 x 169] | Layer-specific intensity rescaling; segmentation and skeletonization; axon and myelination analysis Gamma: 1-5 (Imaris); 0.5 (ImageJ) |
| | | | | MBP | Primary: chicken polyclonal (PA1-10008, Thermo Fisher) Secondary: goat anti-chicken Alexa Fluor 568 (A11041, Thermo Fisher) | | | | | 560 (6 ms; 3.1 mW) | | | | |
| 7 | **Fig. 4 Fig. S12 Movie S5** | **Mouse olfactory bulb (*Npc1*$^{-/-}$) “NPC1”** | BALB/c Npc1$^{nih}$ (*Npc1*$^{-/-}$) | NF-200 | Primary: rabbit polyclonal (N4142-.2ML, Millipore Sigma) Secondary: goat anti-rabbit Alexa Fluor 488 (A11008, Thermo Fisher) | 1.93 (PC-gel) (1x PBS) | Lattice light-sheet/ Light-sheet | 56 x 56 x 135 [108 x 108 x 260] | ~3090 x 5830 x 1850 [~5950 x 11250 x 3570] | 488 (4.5 ms; 1.6 mW) | 405 (100 mW) | 192 for imaging (20 for photodegradation) | Flat-field correction, deconvolution, deskewing/rotation, intensity correction, xy-resampling, stitching, puncta removal Voxel size after deskewing/ rotation (nm): 66 x 66 x 87 [128 x 128 x 169] | Layer-specific intensity rescaling; segmentation and skeletonization; axon and myelination analysis Gamma: 1-5 (Imaris) |
| | | | | MBP | Primary: chicken polyclonal (PA1-10008, Thermo Fisher) Secondary: goat anti-chicken Alexa Fluor 568 (A11041, Thermo Fisher) | | | | | 560 (4.5 ms; 3.1 mW) | | | | |
| 8 | **Fig. S10** | **Human hippocampus slice** | NA | NF-200 | Primary: rabbit polyclonal (N4142-.2ML, Millipore Sigma) Secondary: goat anti-rabbit Alexa Fluor 488 (A11008, Thermo Fisher | 4.40 (PC-gel) (water) | Lattice light-sheet/ Light-sheet | 22 x 22 x 59 [98 x 98 x 260] | ~57 x 52 x 370 [~250 x 230 x 1630] | 488 (10 ms; 0.5 mW) | 405 (100 mW) | 7 for imaging (2 for photodegradation) | Flat-field correction, deconvolution, deskewing/rotation, stitching, puncta removal Voxel size after deskewing/ rotation (nm): 22 x 22 x 22 [98 x 98 x 98] | Gamma: 3.6 (Imaris) |
| | | | | MBP | Primary: chicken polyclonal (PA1-10008, Thermo Fisher) Secondary: goat anti-chicken Alexa Fluor 568 (A11041, Thermo Fisher) | | | | | 560 (10 ms; 1.6 mW) | | | | |
| | | | | Lectin (Binding to glycoproteins) | Lycopersicon esculentum (Tomato) lectin (L-1170-2, Vector Laboratories) conjugated with SeTau-647-NHS (K9-4149, SETA BioMedicals) | | | | | 647 (10 ms; 0.5 mW) | | | | |
| 9 | **Fig. S1** | **Mouse olfactory bulb (wild-type) (serial sectioning)** | BLAB/c (Npc1$^{+/+}$) | NF-200 | Primary: rabbit polyclonal (N4142-.2ML, Millipore Sigma) Secondary: goat anti-rabbit Alexa Fluor 488 (A11008, Thermo Fisher) | 2.00 (Bis-gel) (1x PBS) | Lattice light-sheet/ No photodegradation | 54 x 54 x 130 [108 x 108 x 260] | NA | 488 (6 ms; 1.6 mW) | NA | 47 for imaging gelled and sectioned (total of 3 sections) 69 for imaging sectioned and gelled (total of 3 sections) | Flat-field correction, deconvolution, deskewing/rotation, stitching Voxel size after deskewing/ rotation (nm): 66 x 66 x 87 [128 x 128 x 169] | NA |
| | | | | MBP | Primary: chicken polyclonal (PA1-10008, Thermo Fisher) Secondary: goat anti-chicken Alexa Fluor 568 (A11041, Thermo Fisher) | | | | | 560 (6 ms; 3.1 mW) | | | | |

#### Captions for Supplementary Movies

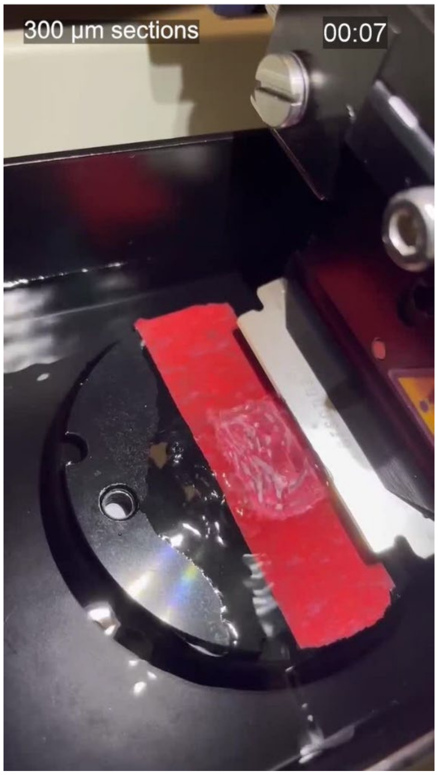


**Movie S1. Vibratome sectioning (~300 µm thick) of a hydrogel-expanded tissue (mouse olfactory bulb) sample in 1x PBS (~2-fold expansion).** The tissue was embedded and expanded in polyacrylamide/sodium polyacrylate gel with bis-acrylamide crosslinker (Bis-gel).

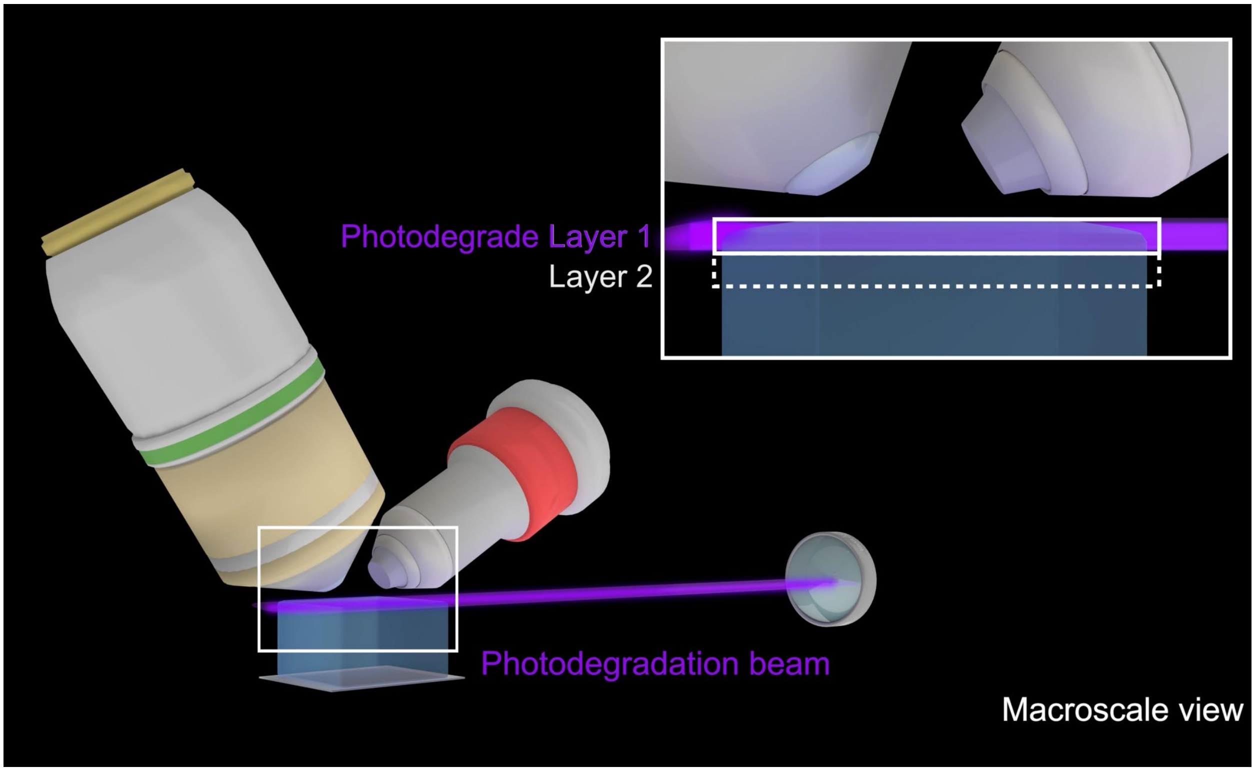


**Movie S2. Animation showing the general concept and workflow of volumetric imaging of biological specimens via photochemical sectioning (VIPS) in both macroscale and nanoscale views.**

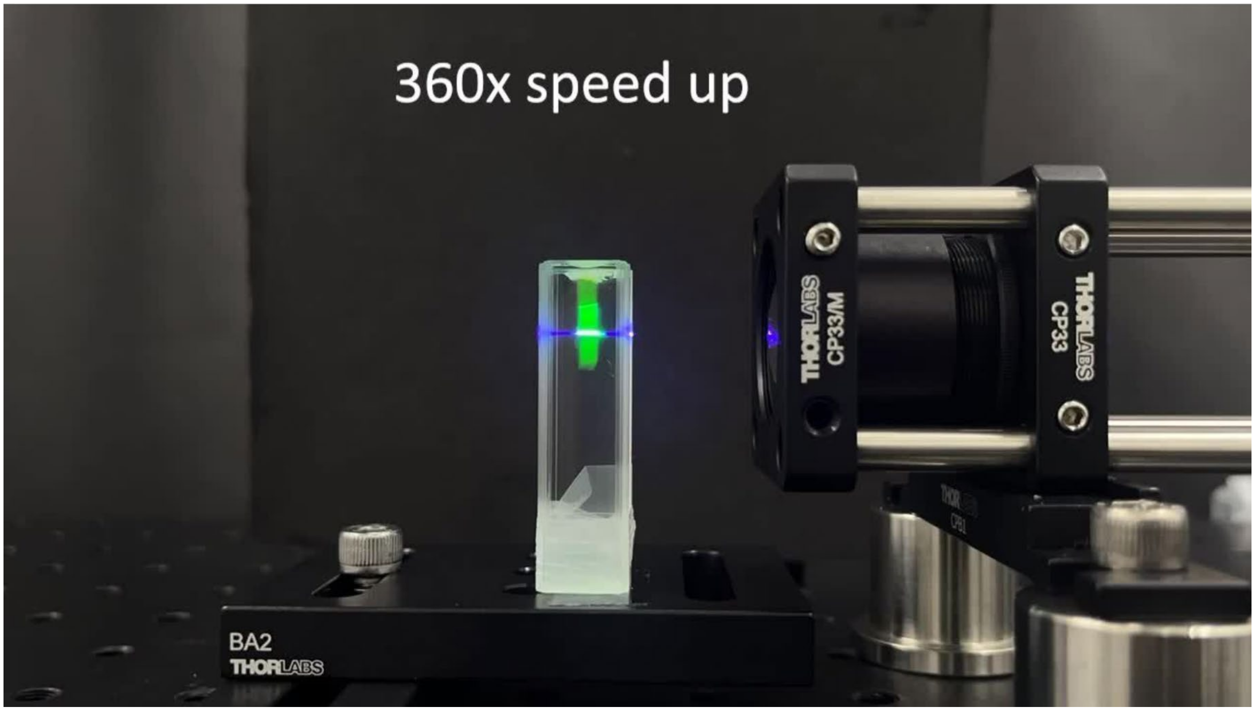


**Movie S3. Spatially controlled photodegradation of photocleavable crosslinker (PC)-crosslinked polyacrylamide/sodium polyacrylate hydrogel (PC-gel).** A piece of fluorescently-labeled PC-gel, hung from the top of a cuvette filled with water, is illuminated with a thin sheet of 405 nm laser from the side.

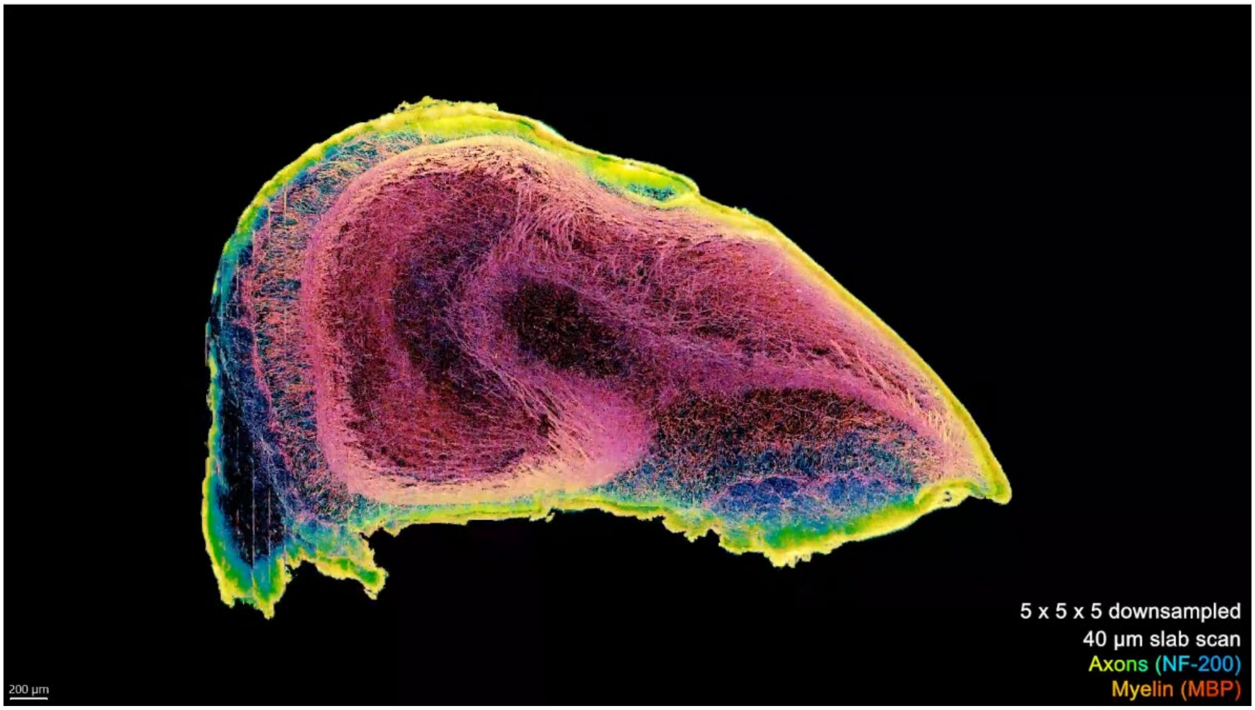


**Movie S4. Volumetric rendering of a 7-week wild-type (WT) mouse OB imaged using sequential lattice light-sheet microscopy and light-sheet photochemical sectioning.** The sample was fluorescently labeled for axons (NF-200) and myelin sheaths (MBP), and expanded ~2-fold using the PC-crosslinked polyacrylamide/sodium polyacrylate hydrogel (PC-gel).

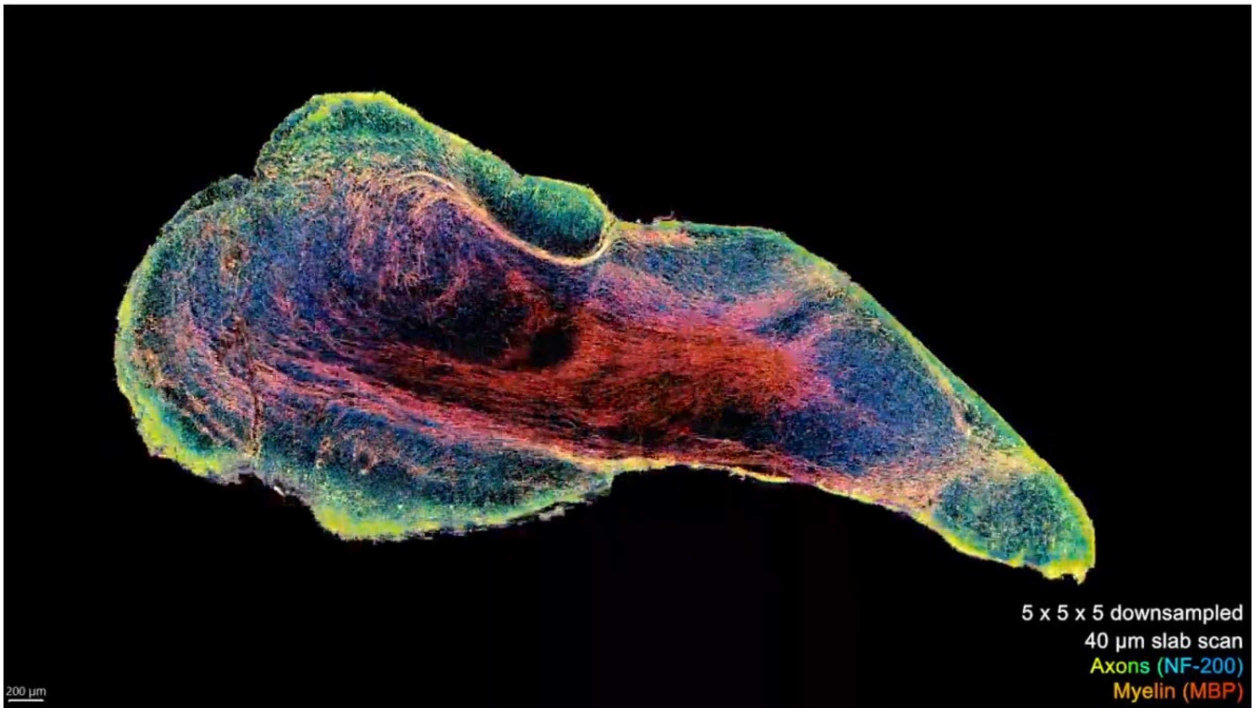


**Movie S5. Volumetric rendering of a 7-week NPC1 (*Npc1*$^{-/-}$) mouse OB imaged using sequential lattice light-sheet microscopy and light-sheet photochemical sectioning.** The sample was fluorescently labeled for axons (NF-200) and myelin sheaths (MBP), and expanded ~2-fold using the PC-crosslinked polyacrylamide/sodium polyacrylate hydrogel (PC-gel).
